# Reconstitution of human DNA licensing and the structural and functional analysis of key intermediates

**DOI:** 10.1101/2024.04.11.589023

**Authors:** Jennifer N. Wells, Vera Leber, Lucy V Edwardes, Shenaz Allyjaun, Matthew Peach, Joshua Tomkins, Antonia Kefala-Stavridi, Sarah V Faull, Ricardo Aramayo, Carolina M. Pestana, Lepakshi Ranjha, Christian Speck

## Abstract

Human DNA licensing initiates the process of replication fork assembly. Specifically, this reaction leads to the loading of hMCM2-7 on DNA, which represents the core of the replicative helicase that unwinds DNA during S-phase. Here, we report the biochemical reconstitution of human DNA licensing using purified proteins, the structural and functional analysis of the process and reveal the impact of cancer-associated mutations on DNA licensing. We showed that the *in vitro* reaction is specific and results in the assembly of high-salt resistant hMCM2-7 double-hexamers, the final product of DNA licensing. We used ATPγS to block complex assembly at the hOrc1-5-Cdc6-Cdt1-MCM2-7 step. We observed that the assembly of this intermediate is independent of hOrc6, although hOrc6 enhances the loading of the second hMCM2-7 hexamer. The structural and mutational analysis of the hOrc1-5-Cdc6-Cdt1-MCM2-7 complex provides insights into hORC-Cdc6 dependent recruitment of hMCM2-7 via five hMcm winged-helix domains. The structure highlights how hOrc1 activates the hCdc6 ATPase, while the analysis of hOrc1 and hCdc6 ATPase mutants uncovered an unexpected role for hCdc6 ATPase in complex disassembly. The structure highlights that Cdc6 binding to Orc1-5 stabilises Orc2-DNA interactions and supports Mcm3-dependent recruitment of MCM2-7. Finally, the structure allowed us to locate cancer-associated mutations at the hCdc6-Mcm3 interface, which showed specific helicase loading defects.

## INTRODUCTION

Prior to cell division, the entire genome needs to be duplicated. This process is split into multiple steps. It starts with loading the replicative helicase onto DNA in the late M early G1 phase, followed by activation of the helicase during the G1/S transition and, finally, assembly of the full replisome and DNA synthesis in S-phase^1, 2^. Helicase loading, also termed pre-replication complex (pre-RC) assembly^3^ and DNA licensing^4^, is a multi-step process^5–7^.

The reaction has been widely studied in budding yeast, where the yeast origin recognition complex (yORC) binds to replication origins in a sequence-specific fashion^8, 9^. In late M-phase, yCdc6 is recruited to yORC^10, 11^, and the yORC-Cdc6 complex encircles DNA^12, 13^. Consequently, and with the help of yCdt1, a spiral-shaped hexameric yMCM2-7 becomes recruited^14, 15^, resulting in a yORC-Cdc6-Cdt1-MCM2-7 (yOCCM) complex^16^. During yOCCM formation, the C-terminal winged helix domains (WHD) of yMcm3 and yMcm7 make the first contact with yORC-Cdc6^16–18^. Then, a yMcm6-Cdt1 interaction promotes the insertion of the DNA into the yMCM2-7 helicase followed by yMCM2-7 closure around DNA^16, 19^ and the open-spiral to closed-ring yMCM2-7 hexamer transition. The induction of ATP-hydrolysis triggers the release of yCdc6 and yCdt1^5, 7^. Subsequently, yORC becomes repositioned to the N-terminal face of the loaded yMCM2-7 hexamer^20^, which promotes another round of yCdc6 and yCdt1-dependent loading of yMCM2-7^21, 22^, resulting in yMCM2-7 double-hexamer formation^23, 24^. This large complex encircles double-stranded DNA and assumes an inactive state. During the G1/S transition, the helicase becomes activated and competent for DNA synthesis^23, 24^. During helicase activation, the MCM2-7 double-hexamer splits and becomes integrated into two replication forks, which replicate DNA bi-directionally. MCM2-7 is the replicative helicase’s motor element within the replication fork, consisting of Cdc45, MCM2-7 and GINS (CMG).

The reconstitution of budding yeast DNA licensing^23, 24^ was instrumental in understanding the process of helicase loading, its regulation and the discovery of the protein structures of the respective intermediates^7, 18–20^. By comparison, human DNA licensing has yet to be reconstituted and is therefore understudied, although the human ORC (hORC) complex has been analysed biochemically and structurally^25–30^. This analysis revealed that hOrc1-5 forms a C-shaped complex similar to yORC. hOrc1, hOrc4, hOrc5 belong to the AAA+ family of ATPases and bind ATP at the subunit interfaces^26, 30^, while hOrc2 and hOrc3 adopt an AAA+ like organisation but do not bind ATP. Finally, hOrc6 adopts a TFIIB-like structure that contains a DNA binding domain^31^. Compared to the yeast complex, hORC adopts a more flexible configuration ^27–29^. In particular, hOrc1 toggles between an ATPase inactive and active state, while the hOrc2-WHD has been observed in two conformations, inactive and autoinhibited, which block hORC-DNA interactions^25, 26, 30^. Interestingly, hOrc6 associates only weakly with hOrc1-5^27–29, 32^, and its role in DNA replication remains ambiguous^31, 33^. Whether the conformational flexibility of hORC has a functional or regulatory role in pre-RC formation is unknown.

hMCM2-7 forms a stable complex in solution that adopts a spiral configuration similar to yMCM2-7, but unlike the yeast complex, does not co-purify with hCdt1^23, 34^. Thus, how hCdt1 becomes recruited during human DNA licensing is not entirely clear. The structure of the hMCM2-7 double-hexamer was recently revealed and identified that base-pairing interactions near the double-hexamer interface are destabilised^35^, which may hinder the sliding of the complex on DNA.

Although some human replication origins have been identified, it is clear that hORC, in contrast to yORC, does not interact with DNA in a sequence-specific fashion. ORC from budding yeast contains an additional alpha helix in yOrc4, an essential contributor to its DNA sequence specificity that is missing in hOrc4^16, 36–38^. Instead, it is thought that the hOrc1 N-terminal bromo-adjacent homology (BAH) domain could be important in chromatin-mediated recruitment of metazoan Orc1-5^39, 40^.

The control of human DNA licensing is crucial. Licensing of DNA that has already been replicated leads to re-replication, promotes recombination and genomic instability^41^. Equally, too little DNA licensing will limit DNA synthesis^42^. In human cells, an excess of hMCM2-7 is loaded, which serve as dormant origins in case an active replication fork becomes terminally stalled^43–46^. However, it is not only the frequency of DNA licensing that is important, but also its speed. Specifically, stem cells require rapid DNA licensing to maintain their pluripotency^47^. Therefore, the process of helicase loading is under the control of the DNA licensing checkpoint that monitors whether the genome has been sufficiently licensed for DNA replication. p53 and RB have an essential role in this checkpoint^48^. Interestingly, cancer cells frequently lose this checkpoint and display greater genomic instability^49^. Therefore, developing clinical therapeutics targeting the DNA licensing machinery represents a promising research avenue^50^.

To date, a number of cancer-associated mutations, as well as several mutations linked to the Meier-Gorlin syndrome, a rare human disease associated with primordial dwarfism^51^, have been mapped to DNA licensing factors. Still, their functional relevance has yet to be discovered due to a lack of structural information and an efficient *in vitro* assay to test the impact of the mutations^52–54^. Here, we report the establishment of a fully reconstituted DNA licensing assay incorporating full-length (FL) proteins, which leads to hMCM2-7 double-hexamer formation. We show that hOrc6, in contrast to yOrc6, is not essential for high-salt stable MCM2-7 loading but improves the efficiency of the reaction. Our data show that hOrc6 is only recruited late during DNA licensing, suggesting that it functions in loading of the second hMCM2-7 hexamer. By blocking ATP-hydrolysis, it is possible to assemble a hOCCM intermediate, which incorporates all human DNA licensing factors with the exception of hOrc6. With cryo-EM, we obtained the structure of the hOCCM, with DNA already inserted into the MCM complex. This structure provides insights into hORC-DNA interaction and allows us to rationalize the structural changes that hOrc1-5 undergoes during hCdc6 binding, which are essential for hMCM2-7 recruitment. Finally, the structure serves as a basis for understanding the impact of cancer-associated mutations on DNA licensing.

## RESULTS AND DISCUSSION

### hOrc1-5 binding to DNA is stabilised by hCdc6

Helicase loading is a multi-step process that hORC-Cdc6 initiates^55^. We started off investigating the ability of hORC and hCdc6 to form a complex and bind to DNA. We focussed on full-length hOrc proteins, as truncations in *Drosophila* Orc1 have been linked to reduced DNA binding^56^ and an N-terminal deletion of hOrc2 has been linked to altered cell cycle profile^57^. The hOrc1-5 complex was overexpressed in budding yeast cells arrested in G1 phase, when S-phase specific cyclin-dependent kinases are inactive, and co-purified as a pentameric complex (Fig. 1a lane 1 and Supplementary Fig. 1a). For the purification, as well as for pull-down assay hOrc1 contained an N-terminal Twin Strep-tag (hOrc1 N-tag). As previous work suggested that hOrc6 does not form a stable complex with hOrc1-5^28^, it was expressed in bacteria and purified as a monomer (Fig. 1a, lane 2 and Supplementary Fig. 1b). hCdc6 was expressed in bacteria and was purified as a homogenous monomer (Fig. 1a, lane 3 and Supplementary Fig. 1b). By employing the Strep-tag on hOrc1 in a pulldown assay, we were able to assess whether hOrc1-5, hOrc6 and hCdc6 would form a complex *in vitro*. We observed that all hOrc1-5 subunits were retained in the hOrc1 N-tag pulldown, while hOrc6 and hCdc6 did not co-precipitate with hOrc1-5 when compared to the non-specific control (Fig. 1b, compare lanes 2 and 3 with 4 and 5). By contrast, when the reaction was repeated in the presence of a 90bp double-stranded (ds) DNA, hCdc6 was retained with hOrc1-5, but not hOrc6 (Fig. 1b, lane 7 and 8). Taken together, these results indicate that neither hOrc1-5-Cdc6 nor hOrc1-5-Orc6 form a stable complex in solution, but rather that the interaction between hOrc1-5-Cdc6 is DNA dependent.

**Fig. 1:**
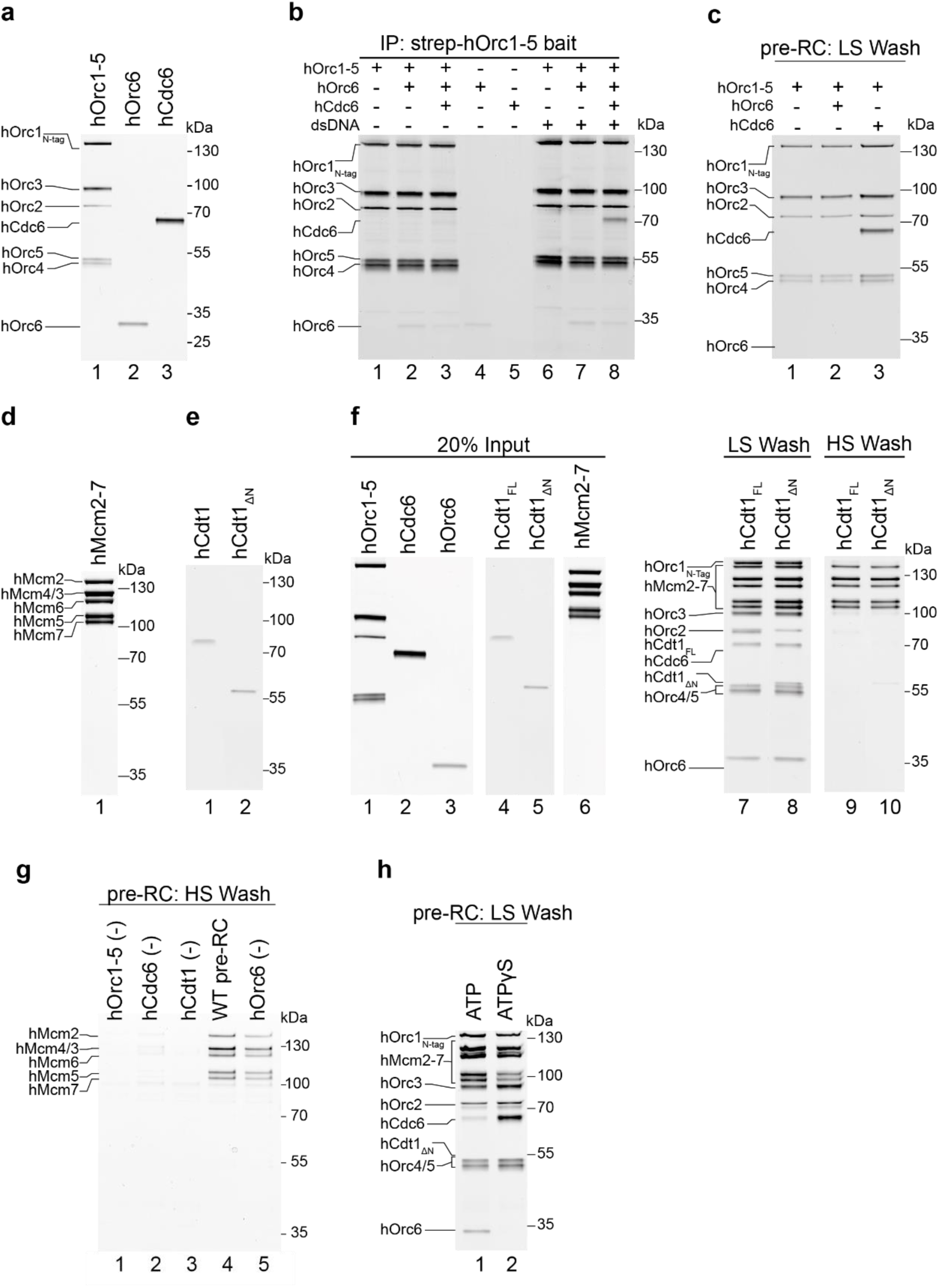
Reconstitution of human DNA licensing using purified proteins. (**a**) Purified hOrc1-5, hOrc6 and hCdc6. (**b**) IP pull down with strep-hOrc1-5 as bait in the presence and absence of a 90bp yeast ARS1 DNA containing the A, B1 and B2 elements. (**c**) Pre-RC assay was performed under low salt conditions with hOrc1-5, hCdc6, and hOrc6. (**d**) Purified hMcm2-7. (**e**) Purified full-length hCdt1 and hCdt1ΔN (aa158-546). (**f**) Pre-RC assay comparing hCdt1 and hCdt1ΔN, in the presence of ATP under low or high salt conditions. (**g**) Specificity of the pre-RC assay under high salt conditions. (**h**) Pre-RC assay under low salt conditions was carried out in the presence of either ATP or ATPγS.

To further validate this observation, we performed an assay with a biotinylated 3 kb human B2-lamin replication origin dsDNA sequence, coupled to streptavidin-coated magnetic beads. Here, protein and DNA were incubated for 20 minutes at 30°C to assess the complex assembly on DNA. While hOrc1-5 only associated with dsDNA alone and only background binding of hOrc6 was observed (compare lane 4 with lanes 2 and 3), enhanced hOrc1-5 binding was observed upon inclusion of hCdc6 (Fig. 1c, lane 1-3). Moreover, comparing ten different 200bp B2-lamin sub-fragments did not show any change in hOrc1-5-DNA interaction, consistent with the concept that hOrc1-5 has no specific sequence specificity (Supplementary Fig. 2a). Taken together, these results indicate that hOrc1-5 bind to DNA in a sequence independent manner, that hOrc6 is dispensable for the initial binding of hOrc1-5-hCdc6 to DNA and that hCdc6 enhances hOrc1-5 binding to DNA.

### Reconstitution of human DNA licensing *in vitro*

Establishing an *in vitro* assay for human DNA licensing has the potential to reveal how the process works at a mechanistic level, enables structural studies and represents a platform to explore the role of patient derived mutations on helicase loading. Full-length hMCM2-7 was expressed in Human Embryonic Kidney (HEK) 293 cells and purified as a hexamer (Fig. 1d and Supplementary Fig. 1d). Full-length (FL) hCdt1 and N-terminally truncated hCdt1_ΔN_ (aa158-546) were expressed in *E*. *coli* and purified as a monomer, though yields for hCdt1_ΔN_ were much higher (Fig. 1e and Supplementary Fig. 1e and f). Consequently, we tested hOrc1-5, hOrc6, hCdc6, hCdt1 or hCdt1_ΔN_, and hMCM2-7 at a relative ratio of 1 : 2 : 2 : 1 : 1.5 respectively (Fig. 1f, lanes 1-6) in the pre-RC assay using a 2kb double-stranded human Lamin B2 replication origin bound to magnetic beads. Here, proteins are assembled on DNA and either washed with low salt buffer preserving reaction intermediates, or, with high salt buffer revealing a salt-stable hMCM2-7 complex that is encircling DNA, which represents the final product of the reaction. After the wash, the DNA-bound proteins are eluted from the beads by nuclease-mediated digest, which ensures that proteins non-specifically associated with the beads are removed. In the low salt wash elution, we observed that all the DNA licensing factors, including hOrc6, were in complex with DNA (Fig. 1f, lanes 7 and 8), and we detected a stable hMCM2-7 loading product in the high-salt washed sample (Fig. 1f, lanes 9 and 10). These results were consistent when testing with both full-length and hCdt1_ΔN_ (Fig 1f, compare lane 7 to 8, and 9 to 10). As yields from hCdt1_ΔN_ purifications were an order of magnitude higher, this construct was predominantly used for subsequent pre-RC assays. To address the specificity of hMCM2-7 loading, we carried out high-salt washed pre-RC reactions in the presence of all factors (Fig. 1g, lane 4), or with individual factors removed (Fig. 1g, lanes 1-3 and 5). In the absence of hOrc1-5, hCdc6 or hCdt1, no high salt stable hMCM2-7 loading was detected. Moreover, to obtain insights into the DNA specificity, we asked whether yeast origin DNA could substitute for human origin DNA. Therefore, we performed comparative low and high salt wash reactions using magnetic beads coupled to either human B2-lamin dsDNA origin sequences or to yeast ARS dsDNA origin sequences. However, no differences in hMCM2-7 loading efficiency were observed (Supplementary Fig. 2b). The data show that high salt stable hMCM2-7 loading is dependent on hOrc1-5, hCdc6, hCdt1 that hOrc1-5, but independent of human origin sequences.

### hOrc6 is supporting high salt stable MCM2-7 loading

Previous work had shown that hOrc6 does not form a stable complex with hOrc1-5 in the presence or absence of DNA^28, 29^. Thus, it was unclear whether hOrc6 participates in human DNA licensing at all, and if so, at which step of the process. Though we did not observe hOrc6 binding in hOrc6-recruitment reactions with hOrc1-5 and hOrc1-5 and hCdc6 (Fig. 1c), we did observe hOrc6 recruitment to hOrc1-5, hCdc6, hCdt1 and hMCM2-7 (Fig. 1f), thus demonstrating that hOrc6 can associate in the presence of all DNA licensing factors. However, the removal of hOrc6 did not block pre-RC formation, but led to a reduction in hMcm2-7 signal in our assay (Fig. 1g, lane 5). Thus, we conclude that Orc6 plays a role in supporting high salt stable MCM2-7 loading. To investigate at which step of the helicase loading reaction hOrc6 associates, we performed complex assembly reactions with ATPγS. This slowly hydrolysable ATP analogue is known to arrest complex assembly at the OCCM stage in budding yeast. Similarly, here we could observe formation of a hOCCM like complex, which is enriched for hCdc6. Interestingly, hOrc6 signal was not observed in the presence of ATPγS, suggesting that hOrc6 predominantly functions during loading of the second MCM2-7 hexamer (Fig. 1h, lane 2).

### Human MCM2-7 forms a salt stable MCM2-7 double-hexamer

It has been observed that chromatin isolated from both yeast and human cells contains high salt resistant MCM2-7 double-hexamers^58, 59^. To understand the relative stability of yeast and human DNA licensing complexes, we assembled pre-RC reactions and then washed them with buffer containing increasing concentrations of sodium chloride. The yeast DNA licensing proteins, yORC, yCdc6, yCdt1 and yMCM2-7 clearly assembled into a complex, which was stable in low salt conditions (Fig. 2a, lane 2). However, after addition of sodium chloride, yORC, yCdc6 and yCdt1 are readily released (Fig. 2a, lane 3). Also, a fraction of yMCM2-7 was retained, and it remained stable when washed with up to 500 mM sodium chloride (Fig. 2a, lanes 3-6). The human DNA licensing proteins assembled into a complex as well (Fig. 2b, lane 2). Upon addition of lower concentrations of sodium chloride, some hORC was retained (Fig. 3b, lanes 3-4), while at high salt concentrations only hMCM2-7 was retained (Fig. 2b, lanes 5-6). Thus, budding yeast and human DNA licensing results in MCM2-7 complexes with nearly identical salt stability. This indicates that DNA licensing results in the topological entrapment of DNA and the establishment of hydrophobic protein-interactions that withstand the high salt wash. We suggest that the inter-hexamer interactions within the DH-interface support the weak protein-protein interactions at the Mcm2-Mcm5 interface^60^.

**Fig. 2:**
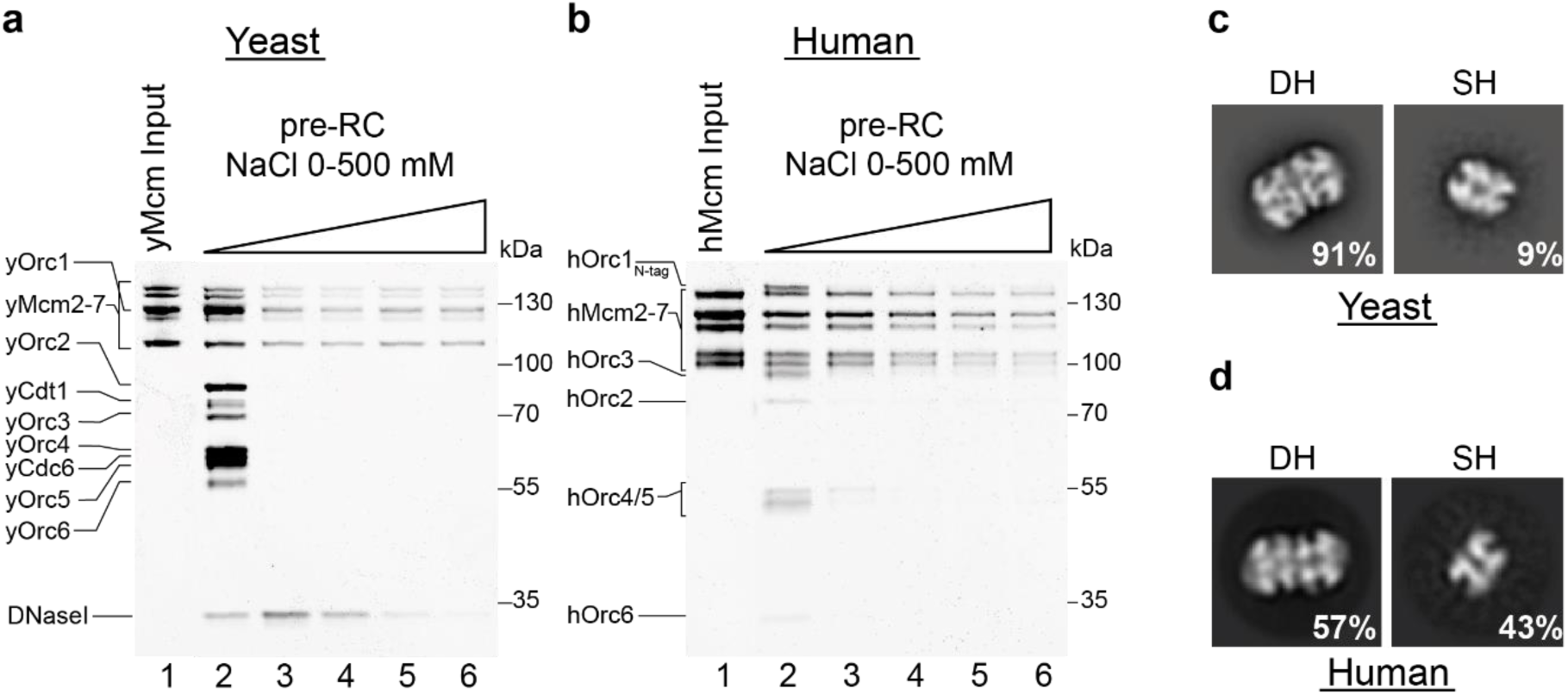
Double hexamer formation with purified yeast and human proteins. Assembled pre-RC reactions were subjected to washes with a sodium chloride gradient ranging from 0 to 500 mM for (**a**) yeast and (**b**) human. Negative stain EM 2D class averages generated with CryoSPARC carried out on pre-RC assay reactions washed with 300 mM NaCl from (**c**) yeast and (**d**) human. The percentage numbers represent the proportion of the total particles in each class of either double hexamer (DH) or single hexamer (SH). hMCM2-7 DH formation was more variable than yMCM2-7 DH-formation, suggesting additional regulation mechanisms exist.

**Fig. 3:**
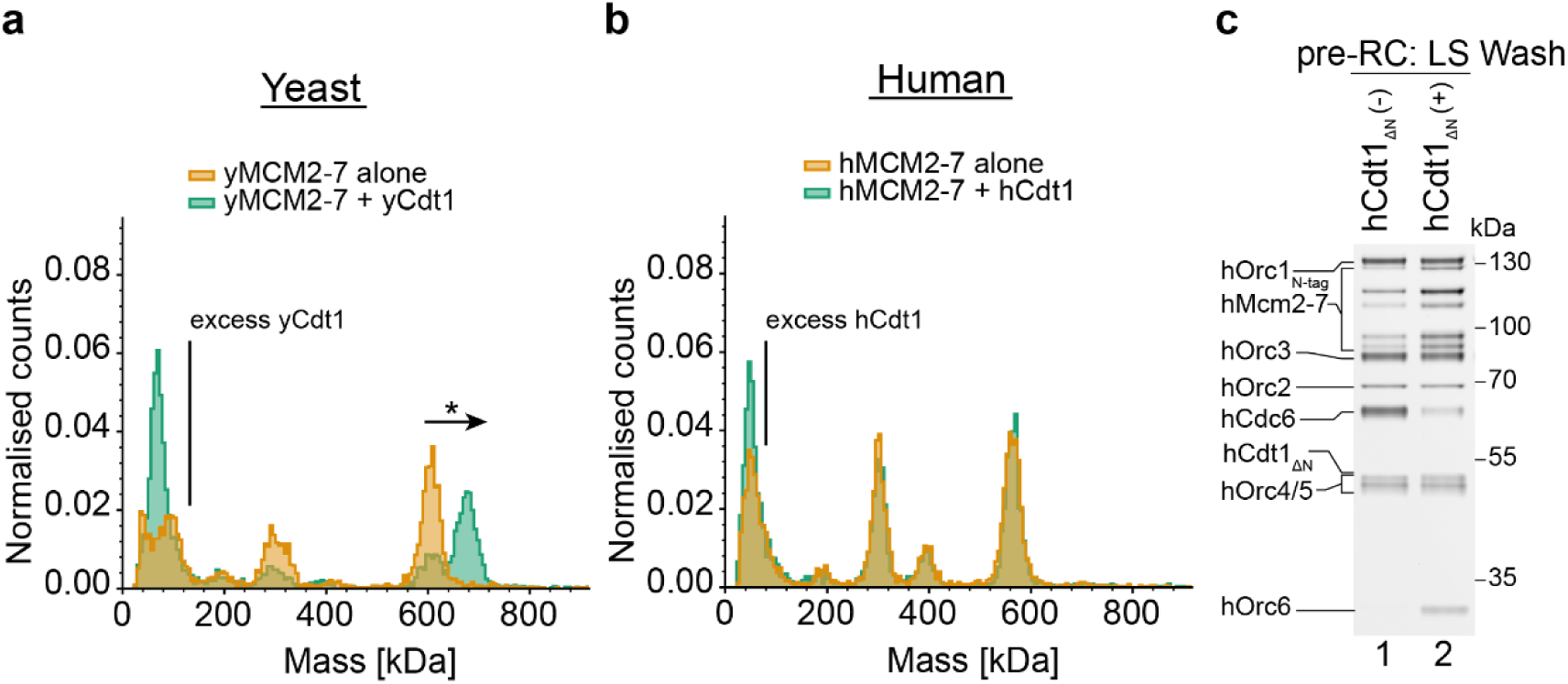
hCdt1 and MCM2-7 do not interact in solution, but hCdt1 promotes hMCM2-7 recruitment during pre-RC formation. Mass photometry on (**a**) yeast and (**b**) human hMCM2-7 and hCdt1 was carried out in solution with a 2-fold excess of hCdt1. (**c**) Pre-RC assay under low salt conditions was carried out in the absence and presence of hCdt1ΔN, highlighting that initial hMCM2-7 recruitment occurs without Cdt1.

Next, we wanted to ask whether the final human helicase loading product assumes the MCM2-7 double-hexamer, as previously observed with the budding yeast reaction^23, 24, 61^. Negative stain electron microscopy was applied to eluates from the high salt washed pre-RC reactions carried out using both the human and yeast systems. The 2D class averages confirm formation of MCM2-7 double hexamer for yMCM2-7 (Fig. 2c) and for hMCM2-7 (Fig. 2d). Interestingly, in context of human DNA licensing, a larger fraction of single-MCM2-7 was retained. This could be due to reduced hMCM2-7 double hexamer stability, or could suggest that additional protein factors or specific DNA sequences may regulate the reaction. Future work has the potential to reveal how human DNA licensing can be regulated by post-translational modifications or chaperones, which could change the reaction dynamics. In summary, this data shows that the human DNA licensing assay supports high salt stable, hMCM2-7 complex formation on DNA, and the assembly of MCM2-7 double hexamers, which represent the end product of the reaction. Thus, the human DNA licensing assay is fully functional and well suited for structural and functional analysis.

### The role of hCdt1 in hMCM2-7 recruitment

In budding yeast, yMCM2-7 and yCdt1 form a complex, which stabilizes the spiral-shaped MCM2-7 complex^14^. In contrast, hMCM2-7 purified from human cells does not co-purify with hCdt1^34^. However, hCdt1 was found to interact with hMcm6 in pulldown assays^62^. Therefore, we used mass-photometry to analyse whether hCdt1 and hMCM2-7 interact in solution, using homologous yeast proteins as a positive control. We observed that the addition of yCdt1 to yMCM2-7 resulted in a yCdt1-MCM2-7 complex, identifiable by an upshift in the maximum molecular mass detected (Fig. 3a), but this was not true for human homologues (Fig. 3b). The same results were obtained using hCdt1_FL_ and hCdt1_ΔN_ (Supplementary Fig. 2c and 2d). This result raised the question of whether hCdt1 is required for the initial binding of hMCM2-7. To address this question, we performed pre-RC assembly reactions in the absence and presence of hCdt1_ΔN_. When hCdt1_ΔN_ was omitted, we observed, to our surprise, association of all Mcm subunits, although at reduced levels (Fig. 3c, lane 1). Thus, the data suggest that hCdt1 and hMCM2-7 do not form a complex in solution but act synergistically during pre-RC formation to promote hMCM2-7 recruitment.

### Cryo-EM analysis of the human OCCM

Many of the proteins involved in human helicase loading belong to the AAA+ ATPase family, specifically hOrc1, hOrc4, hCdc6 and hMCM2-7. Thus, the slowly hydrolysable ATP analogue, ATPγS, can be used to enrich for short-lived early helicase loading intermediates (Fig. 1h). We performed pre-RC assays with ATPγS and analysed the obtained complexes by negative stain electron microscopy. The resulting 2D class averages are consistent with known reaction intermediates including hMCM2-7 single hexamers, the hOCCM intermediate, and fully loaded double hexamers (Fig. 4a). Nearly half of the particles could be attributed to the OCCM (Supplementary Fig. 3), characterised by a smaller ring corresponding to ORC-Cdc6 which sits atop a larger Mcm hexamer. As such, the data demonstrate that complex formation was enriched at the OCCM step, and that ATP hydrolysis is important for release of the helicase loader from the helicase.

**Fig. 4:**
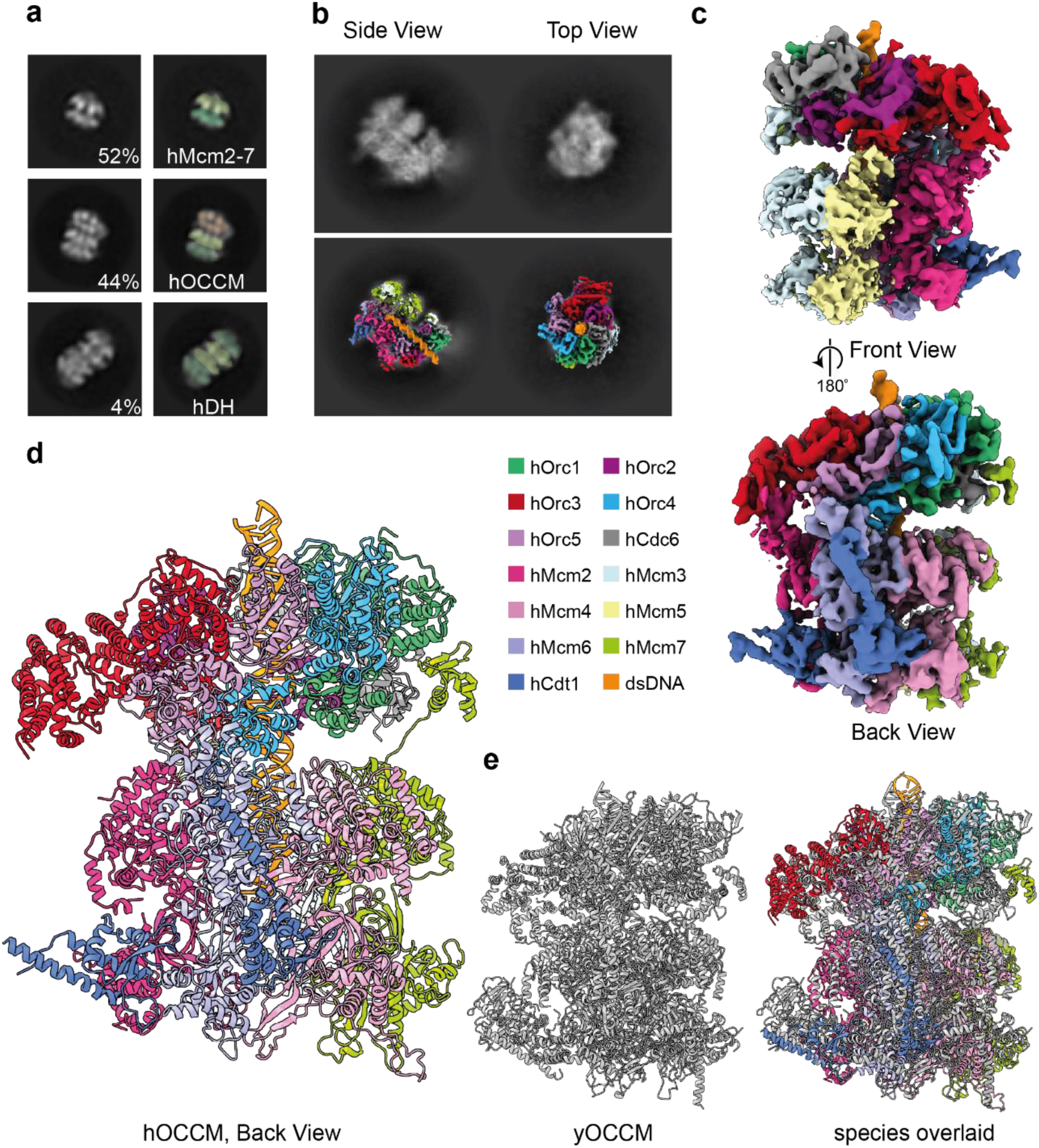
The electron microscopy structure of the hOCCM. (**a**) Negative stain class averages of a low-salt washed pre-RC reaction containing ATPyS. We observed the following pre-RC complexes: single hMCM2-7 hexamers, hOCCM and hMCM2-7 double hexamers. The green densities represent hMcm rings and the pink densities hORC-Cdc6 rings. (**b**) Representative 2D class averages of hOCCM top and side views are in the upper part, and projections of the 3D model on the 2D class are in the lower part. (**c**) The segmented experimental density map is shown from different angles and coloured by protein subunit. (**d**) The hOCCM molecular model was built from the cryo-EM structure (PDB: 8RWV). (**e**) Structure comparison of hOCCM (in colours) with yOCCM (grey, PDB: 5V8F).

Next, we aimed to structurally resolve the hOCCM by cryo-EM. The sample was assembled and consequently grids were prepared. The data were collected on a Titan Krios 300kV microscope and processed using the CryoSPARC2 software suite^63^. The representative 2D class averages are consistent with a hOCCM complex (Fig. 4b). The final 3D reconstruction from 8,730 particles resulted in a structure with an average overall resolution of 6.09 Å (Fig. 4c, Supplementary Fig. 3 and 4, Supplementary Table 1). This allowed for building of a 3D molecular model with good confidence in backbone trajectories (Fig. 4d). As shown in the local resolution analysis, the upper tier consisting of ORC-Cdc6 is adopting a more fixed conformation, while the lower tier consisting of Cdt1-MCM2-7 is more flexible (Supplementary Fig. 3e). As expected, the comparison of the hOCCM and yOCCM indicates that the overall organisation is similar (Fig. 4e, Supplementary Figure 5a, b). Upon closer inspection, we can identify a number of important differences, which start to explain how human DNA licensing differs from yeast DNA licensing.

### hMCM2-7 encircles DNA and adopts an open ring structure

Within the pre-RC intermediate, hMCM adopts a partially closed ring conformation. The DNA is observed inside the central hMCM2-7 channel; thus, the helicase has already been loaded on DNA (Fig. 5b, d). As in the yeast OCCM^16^, the DNA is not visible within the N-terminal Mcm2-7 ring section (Fig. 5d), likely due to DNase mediated digestion. Consistently, we observe that 38bp of dsDNA running through the core of the hOCCM has a similar length as the 39bp found in the yOCCM structure^16^. The hMCM2-7 ring is partially opened at the Mcm2-Mcm5 gate, and the hMcm5 density is not well resolved, consistent with high local flexibility (Fig. 5c, Supplementary Fig. 3e and 5a). Large sections of hCdt1 can be seen in the structure, with the N-terminal section of hCdt1 binding to the hMcm2/6 interface and the C-terminal section to hMcm6/hMcm4, its position within the hOCCM seemingly similar to yCdt1 (Fig. 5d, Supplementary Fig. 3g and 5b). The two domains of hCdt1 are connected by a flexible loop region, that was not resolved. Although Cdt1 diverged significantly during evolution, with only 10.7% identity and 17.1% similarity between yeast and human Cdt1 amino acid sequences, the overall 3D structure and positioning of hCdt1 is conserved from yeast to human.

**Fig. 5:**
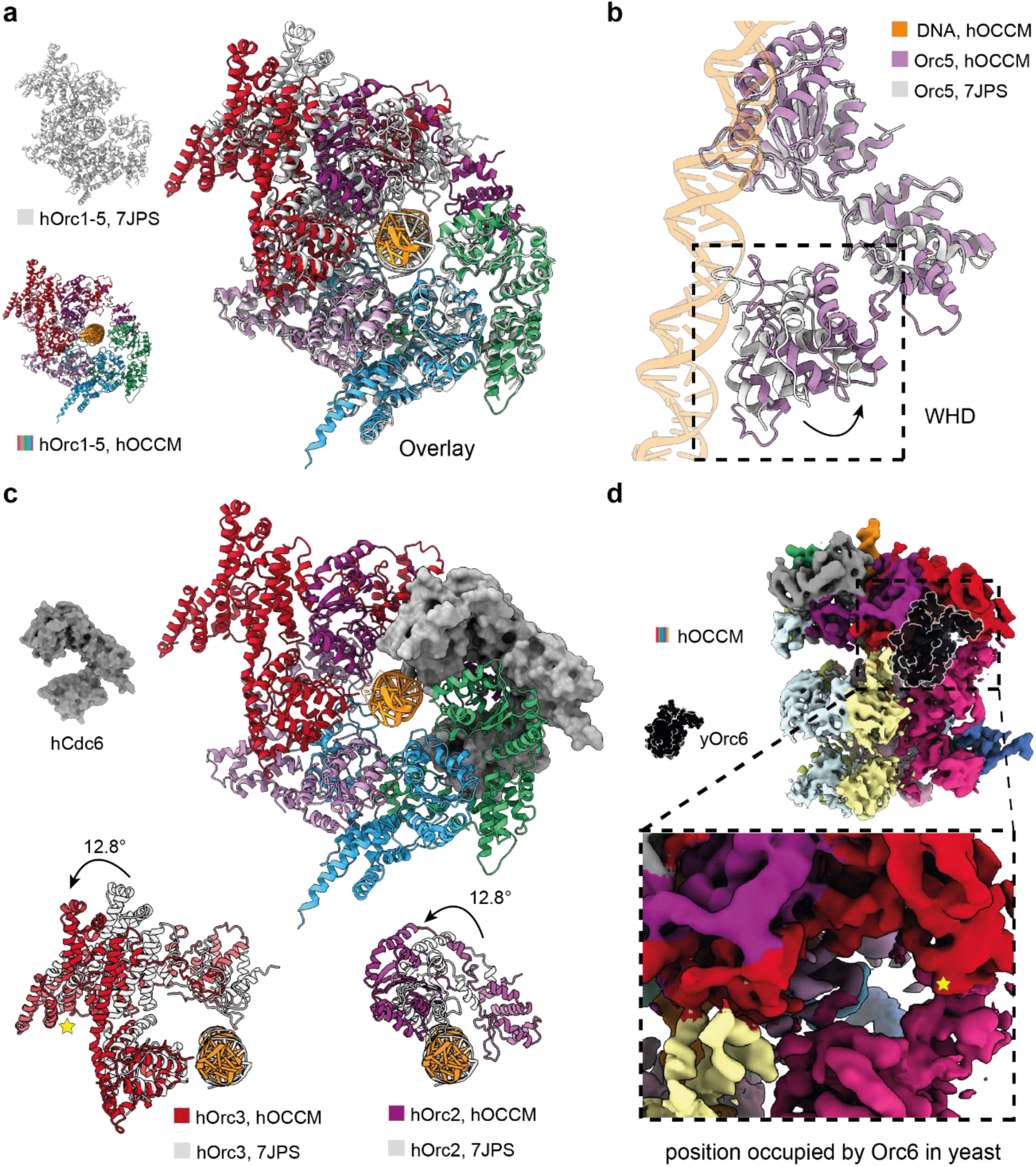
hORC-Cdc6 in the hOCCM structure. (**a**) Comparison of hOCCM (coloured) with human hOrc1-5-DNA structure (grey, PDB: 7JPS). (**b**) Reorganisation of hOrc5 WHD in hOCCM when compared to hOrc1-5-DNA structure (grey, PDB: 7JPS). (**c**) hOrc2-Orc3 become reorganised to accommodate hCdc6, displayed with surface rendering, in hOCCM compared with to hOrc1-5-DNA structure (grey, PDB: 7JPS). (**d**) No densities were observed in the position occupied by yOrc6 in yOCCM (black, PDB: 6WGG). A star marks the establishment of new contact between hMcm2-WHD and Orc3 in the hOCCM structure.

### Organisation of hOrc1-5-hCdc6 within the hOCCM

In the hOCCM complex, hORC-hCdc6 adopt a ring-shaped organisation with an upper tier made up from the AAA+/AAA+-like domains and a lower tier made up of the WHDs, with DNA located in the centre. In order to compare the organisation of hORC in the hOCCM to the hOrc1-5-DNA structure (PDB: 7JPS), we aligned both structures via hOrc1. This highlights that the motor module consisting of hOrc1-hOrc4-hOrc5 adopts a similar overall conformation (Fig. 5a). The largest changes in the motor module are observed within hOrc5-WHD, which is repositioned in the hOrc1-5-DNA structure, likely reflecting that only a very short 13bp DNA could be observed in the published hOrc1-5-DNA complex (Fig. 5b)^30^. Interestingly, more substantial changes can be observed outside of the hOrc1-hOrc4-hOrc5 motor module. Specifically, Orc2 and Orc3 are rotated by 12.8°, which generates a gap between Orc1 and Orc2, that accommodates hCdc6 (Fig. 5c).

We compared available structures from yeast and *Drosophila* in order to localise where hOrc6 would dock to hOrc3, based on predicted interaction surface^16, 64^. We note that while the predicted hOrc6 binding surface in hOrc3 is accessible, no hOrc6 was observed (Fig. 5d). As such, our structural data are consistent with our biochemical observations which showed a lack of hOrc6 binding at the hOCCM stage (Fig. 4a and 4c), which is markedly different from the yeast and *Drosophila* counterparts^16, 64^. We speculate that human Orc6 could be recruited in an alternative fashion to the pre-RC, likely involving an interaction with the loaded MCM2-7 hexamer. Indeed, the N-terminal domain of hMcm6 has been predicted to interact with hOrc6 ^65^, very similarly as the yMcm5 interaction with yOrc6^20^.

### Role of ATP-hydrolysis in human pre-RC formation

Inspection of the interface of hOrc1 and hCdc6 revealed that a loop in hOrc1, which has been unstructured in the context of the hOrc1-5-DNA structure^30^, became structured within the hOCCM and formed a short helix (Fig. 6a). This hOrc1 helix contains the conserved arginine finger, which we observed to be proximal to the Walker B motif of hCdc6. It is well established that these two motifs form in AAA+ proteins a composite ATPase site^66^. Thus, the hOCCM structure suggest that hOrc1 and hCdc6 form upon complex formation an active ATPase pair. Moreover, hOrc1 and hOrc4 also form a composite ATPase pair, with a conserved arginine finger in hOrc4 and a Walker B motif in hOrc1. However, when comparing hOrc1-5-DNA with the hOCCM, we did not observe any structural change for the hOrc1-Orc4 ATPase pair.

**Fig. 6:**
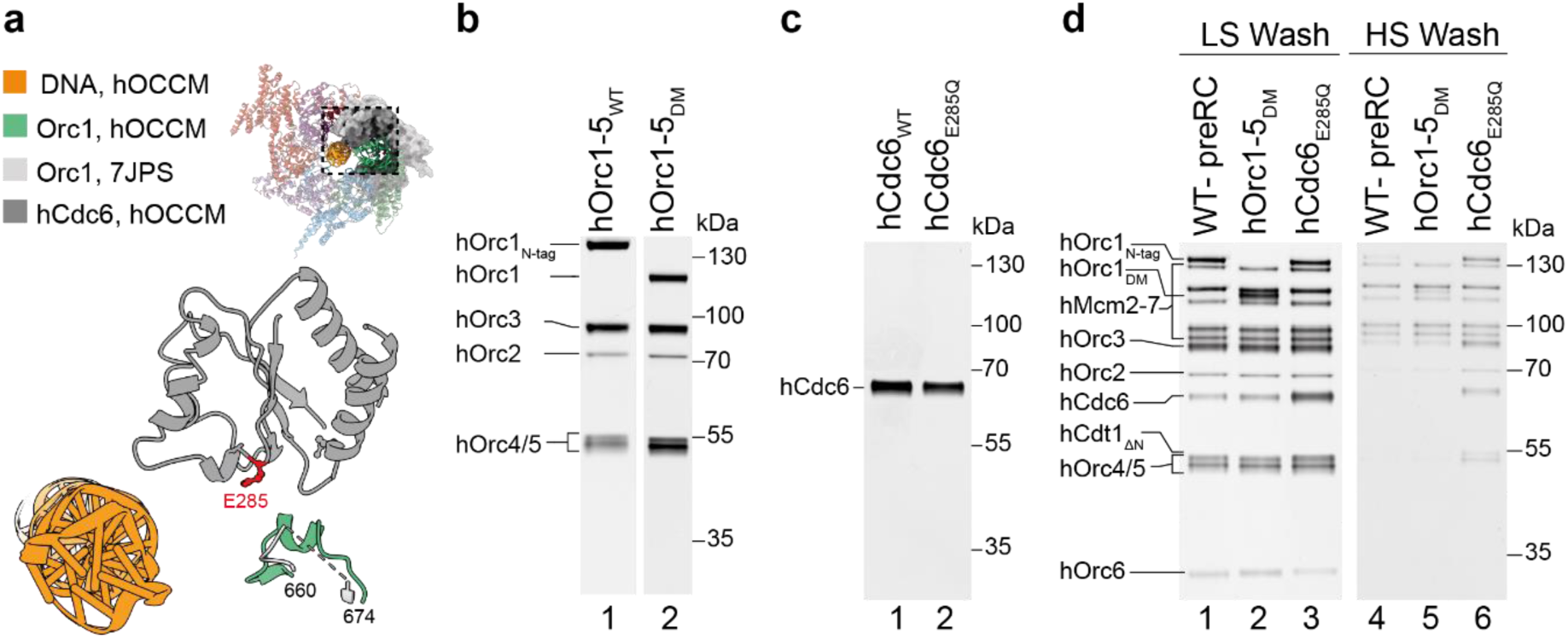
Cdc6 becomes restructured in the OCCM and is crucial for complex disassembly. (**a**) Stabilisation of an hOrc1 helix formed by residues 665-670, which is unstructured before hCdc6 binding as evidenced by docking hOrc1 (PDB: 7JPS) in light grey and Orc1 of hOCCM in green. (**b**) Purified hOrc1-5 and hORC1-5_DM_ (Orc1E621Q/hOrc4R209A). (**c**) Purified hCdc6 WT and hCdc6E285Q. (**d**) The pre-RC assay was carried out under low and high salt wash conditions with WT Orc1-5, Orc1-5 DM or Cdc6 E285Q.

We wanted to investigate whether the observed hOrc1 or hCdc6 ATPases function in human pre-RC formation. To that end, we produced an hOrc1 ATPase mutant, which combines the hOrc4 R209A arginine finger mutation and the hOrc1 E621Q Walker B mutation (Fig. 6b, lane 2), both of which effect hOrc1 ATP-hydrolysis. We also generated in Cdc6 the E285Q Walker B ATPase motif mutation (Fig. 6c, lane 2). Both mutants were tested in the pre-RC assay: we observed that in the presence of low salt, complex assembly occurred with comparable recruitment of Orc1-5 and MCM2-7 (Fig. 6d, lanes 1-3). The even recruitment of hOrc6 across the wild-type (WT) proteins and the mutants indicates that complex formation proceeded past the OCCM stage, even in the presence of the ATP hydrolysis deficient hOrc1-5 and hCdc6. We next observed that the Orc1 double mutant supported high-salt stable MCM2-7 complexes (Fig. 6d, lane 5). Thus, the composite hOrc4-hOrc1 ATPase interface seemingly does not play a major role in DNA licensing. Interestingly, in the presence of Cdc6 E285Q, ORC-Cdc6 were also stabilised on DNA under high salt wash conditions, highlighting that in fact Cdc6 ATP-hydrolysis is important for ORC-Cdc6 disassembly (Fig.6d, lane 6). Crucially, in budding yeast none of the yORC, yCdc6 or yMCM2-7 ATPase mutants have led to a stabilisation under high salt conditions^67^. However, an *in vivo* analysis of a yCdc6 ATPase mutant revealed that the mutant supported MCM2-7 binding to origins, but cells remained in S-phase. Interesting, microinjection of an hCdc6 ATPase mutant in human cells in G1 also inhibited DNA replication ^68^. As such, these results reveal that hCdc6 ATP-hydrolysis plays an essential role in human pre-RC disassembly, a finding that is consistent with previous cellular experiments in yeast and human.

### hORC-hCdc6-DNA interactions in the OCCM complex

The hOCCM structure provides important insights into hOrc1-5-hCdc6-DNA interactions, as the DNA extends through the entire hORC-hCdc6 complex. 38 bp of the DNA in the cryo-EM densities were modelled (Supplementary Fig. 3f). When comparing the yeast and human OCCM, it is clear that in yeast, the DNA is bent towards yOrc2 ISM by 10°, while the human DNA is comparatively straight (Fig. 7a). As in the yOCCM, the DNA in the hOCCM was deformed from the standard B-form, with a widening of the major groove and a narrower minor groove (Fig. 7a). While the resolution was limiting for the DNA in the hOrc1-5-DNA structure^30^, we observed a clear density for the DNA in the hOCCM experimental map (Supplementary Fig. 3f). These results indicate that while hORC is not known to bind to DNA in a sequence-specific manner, it recognizes DNA that is either naturally deformed, or the complex itself introduces structural changes in the DNA upon binding, similar to what has been observed for yORC^36^.

**Fig. 7:**
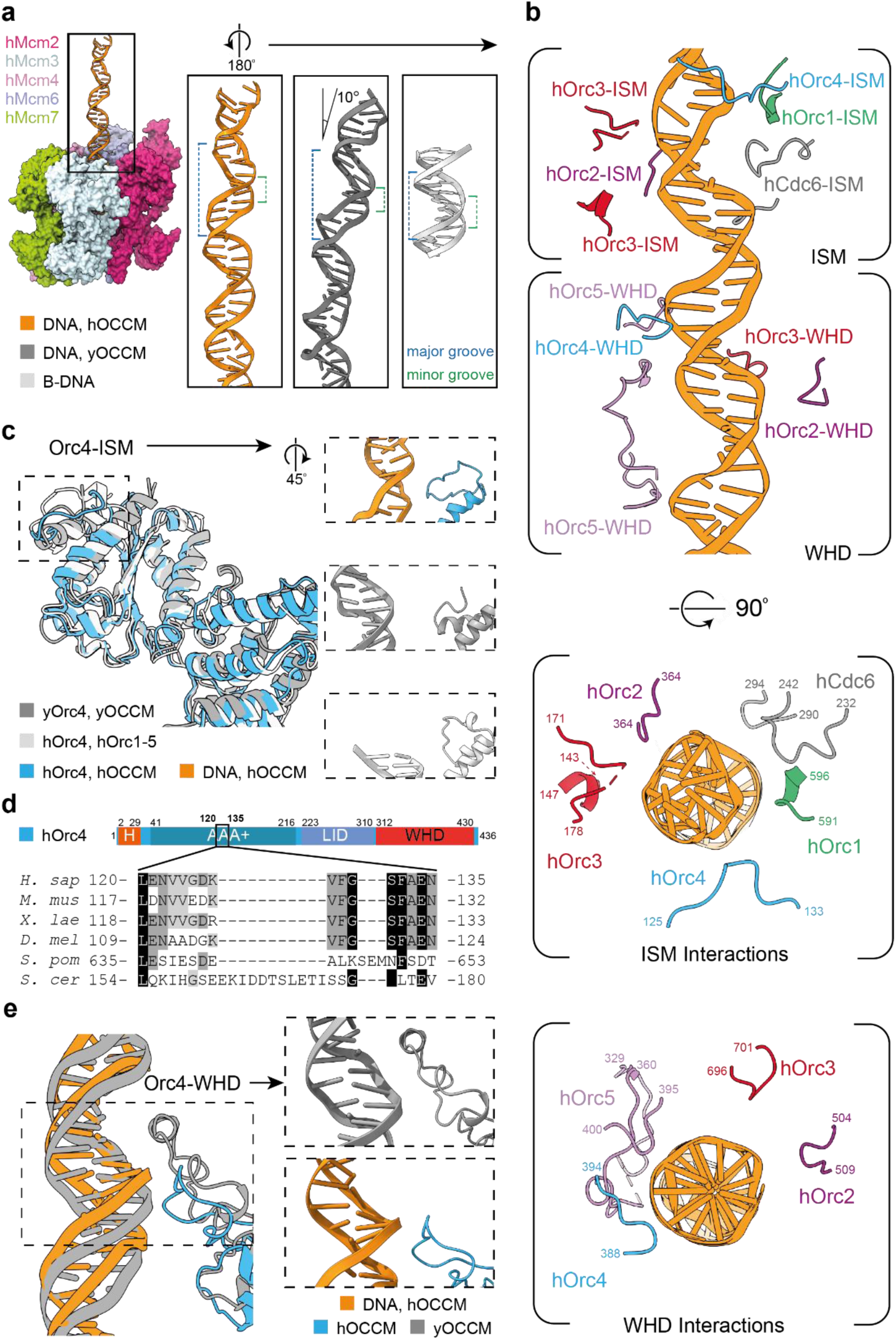
DNA structure and interactions in hOCCM. (**a**) The DNA interacting with yORC1-5 in the yOCCM (PDB: 58VF; grey) is bent by 10° respect to that of the hOCCM (orange). The hOCCM DNA is stretched with respect to the canonical B-form DNA (light grey) in the upper part of the DNA. (**b**) Detailed hOrc1-5 and hCdc6 interactions with DNA. The upper deformed part of the DNA is contacted by the ISM, and the lower less deformed part by WHDs. The view is rotated 180° with respect to figure (a). (**c**) hOrc4-DNA interactions. hOrc4 (light blue) is positioned closer to the DNA in hOCCM than in yOCCM (PDB: 58VF; grey). The DNA is not well-resolved enough in hOrc1-5 (PDB: 7JPS; light grey). (**d**) hOrc4 sequence alignment. The ISM loop region is longer in yeast containing a helix insert, while higher eukaryotes contain a conserved alternative ISM sequence. (**e**) Alternative Orc4 WHD close interactions with DNA in the human (blue) and yeast (grey) proteins.

Beyond these changes in the DNA structure, in the hOCCM we also observed a series of contacts between ORC proteins and the DNA backbone. Specifically, the hOrc and hCdc6 subunits interact with the DNA via two different modules: 1) the initiation-specific motifs (ISMs) in the AAA+ domains and 2) the hairpin and inserted helix in the WHDs (Fig. 7b). Reflecting the ‘J’ shaped tertiary structure of the individual proteins, ISM contacts localise to the upper part of DNA, toward the N terminus of hOrc1-5/hCdc6, while the WHDs exclusively contact DNA more buried in the complex, towards the C terminus of hOrc1-5/hCdc6, which forms interactions with hMCM. Interestingly, the aforementioned structural deformations to the DNA (Fig. 7a) dominantly localise to the more superficial region of DNA that is forming contacts with the ISMs of hOrc1-5/hCdc6. As such, the ISMs are responsible for either reading out this structural change, naturally occurring in the DNA, or directly inducing these changes in the structure of the DNA. Upon comparison with the hOrc1-5-DNA structure (PDB: 7JPS)^30^, we observed additional protein-DNA interactions in the hOCCM. This improved resolution likely reflects an increase in the stability of the DNA in hOCCM, where the length of the DNA is significantly longer.

We observed that the hOrc1-ISM tracked the phosphate backbone via a loop formed by residues 591-596, in agreement with observations reported from the hOrc1-5-DNA structure (PDB: 7JPS) (Fig. 7b)^30^. By contrast, we observe that the hOrc2-ISM, located within the loop spanning residues 364-368 is shifted into the direction of hOrc3, and positioned within range to facilitate phosphate-backbone contacts with the DNA. This shift represents a difference from the position adopted by the hOrc2-ISM loop in the hOrc1-5-DNA structure. Moreover, in hOCCM the hOrc2-WHD, which was not visible in the hOrc1-5-DNA structure, is stabilised and hovers in proximity to the DNA without making direct contacts (Fig. 7b).

Despite discontinuous density in the flexible loop regions of hOrc3, a clear interaction is observed between the phosphate-backbones of the DNA and a positively charged ISM loop spanning residues 171-178 (Fig. 7b). A downstream helix formed by hOrc3 residues 143-147, also of the ISM, is slightly more distal to DNA, as is the hOrc3-WHD residues 696-701 (Fig. 7b). In general, hOrc3 contacts appear closer in the context of the hOrc1-5 structure than the hOCCM.

hOrc4 makes two contacts with the DNA backbone. The hOrc4-ISM contains a loop (residues 125-133) that directly binds DNA, differing from the yOCCM and hORC-DNA structures where the loop was oriented more distally to the DNA (Fig. 7c). A sequence alignment shows that the ISM loop region diverged in higher eukaryotes, reducing a longer loop found in budding yeast (Fig. 7d). Moreover, the hOrc4-WHD makes a very close contact with the DNA. Importantly, the hOrc4-WHD does not contain the insertion helix which in yOrc4 confers specificity to DNA sequence motifs. Instead, a short loop formed by residues 388-394 of hOrc4 interacts with the DNA minor groove, rather than the major groove as for the yOrc4 insertion helix. This human Orc4-WHD loop is well positioned to make phosphate backbone contacts with the DNA, which in yeast is too distal to engage (Fig. 7e). These observed DNA contacts appear closer in the context of the hOCCM than the Orc1-5-DNA complex, which may reflect that the protein is able to form more stable contacts than could be observed using shorter DNA fragments.

The hOrc5-ISM is not in contact with DNA, but the hOrc5-WHD makes three contacts with the DNA (Fig. 6b, 7b, Supplementary Fig. 6a). hOrc5 tracks both phosphate backbones of a minor groove by a short loop spanning aa395-400 and aa353-358. The latter forms an extended loop spanning residues 329-360, where Orc5 aa335-339 makes a separate, third contact with the phosphate backbone. Again, more extended hOrc5 DNA contacts have been observed within the hOCCM than in the hOrc1-5-DNA complex.

The shorter, partially resolved hOrc5 loop spanning residues 395-400 is conserved in both pairwise sequence alignments and molecular localisation with yOrc5 (aa 436-450) (Supplementary Fig. 6b, 6c). Considering that this loop forms stabilising interactions with both the AAA+ and WHDs of hOrc2 in the autoinhibited hOrc2-5 complex^25^ (Supplementary Fig. 6c), and that the corresponding loop in yOrc5 is crucial in the context of DNA bending^36^, this patch may have a regulatory role in DNA licensing.

Notably, we observe residues 335-339 of the long hOrc5 loop engaging a region of the DNA nearing the N-terminal of Mcm, which likely only become structured as phosphate backbone contacts are established in the OCCM structure. Intriguingly, a basic patch formed by this region, conserved in higher eukaryotes, is not observed in sequence alignments or molecular localisation in yeast (Supplementary Fig. 6d, e). Unfortunately, our attempts to replace charged amino acids with alanines in this region resulted in destabilisation of the protein complex.

Finally, the hCdc6-ISM makes multiple contacts with the backbone of DNA (Fig 7b). The hCdc6-WHD is too distant to contact the DNA, owing to a shorter wing in contrast to the yCdc6 WHD. Considering that hCdc6 stabilizes Orc1-5 on DNA (Fig. 1c), it possible that these contacts and the Cdc6 induced reorganisation of hOrc2 and hOrc3 result in the improved DNA binding. Still, the *in vivo* relevance of this improved DNA interaction remains to be established.

### Five MCM WHDs stabilise the ORC-MCM interface

At the hORC-MCM interface, we observed density for five Mcm C-terminal WHDs, contrasting to only four WHDs which have been reported engaging with ORC in the yOCCM (Fig. 8a).

**Fig. 8:**
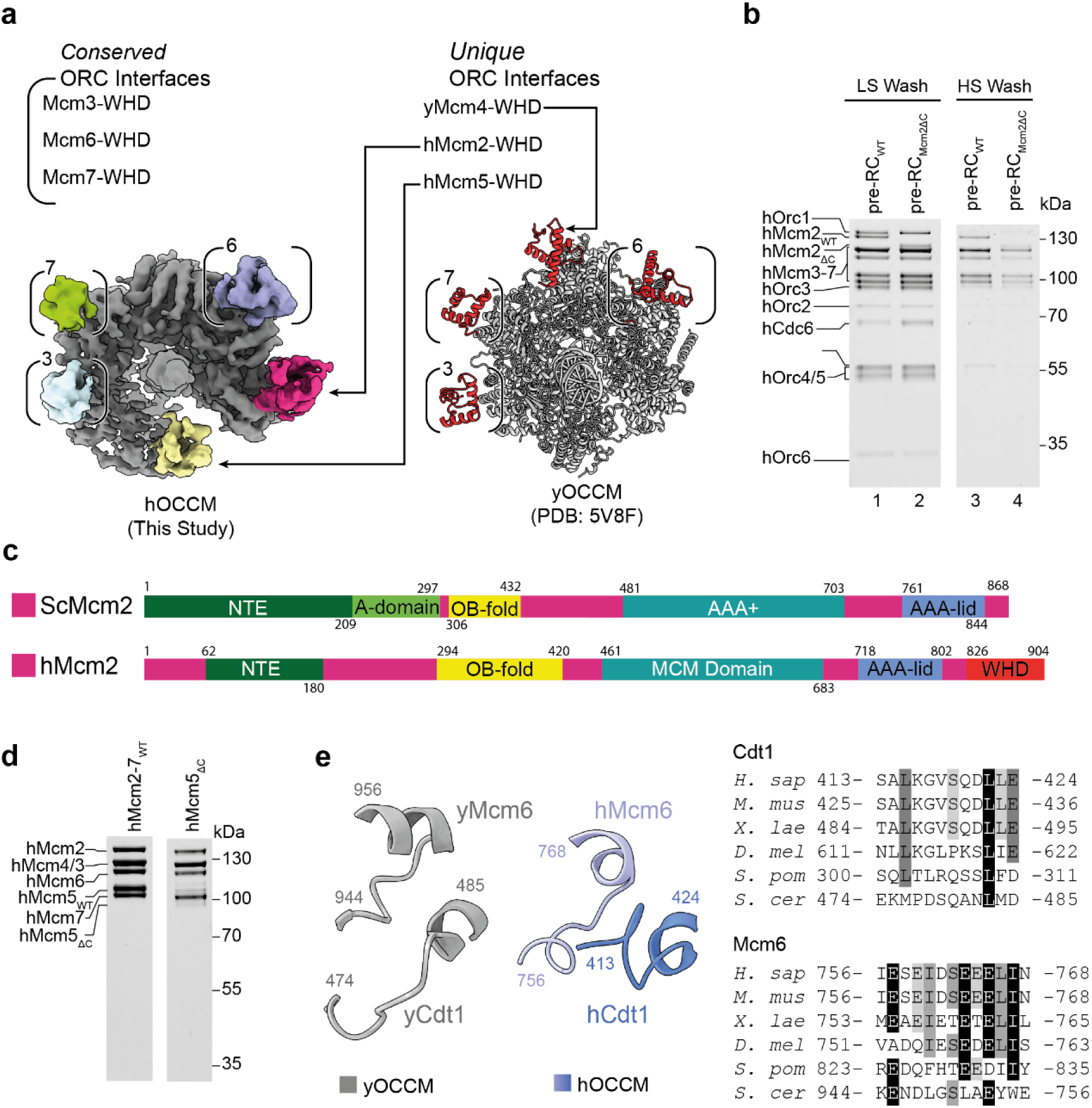
Five hMCM2-7 WHDs make contact with hOrc1-5-Cdc6. (**a**) Mcm WHDs (coloured) contacting Orc-Cdc6 (grey) in hOOCM and yOCCM (PDB: 5V8F). The conserved interfaced are highlighted in brackets (top left), and the unique interfaces in the top right. (**b**) Pre-RC assay under low and high salt wash conditions was carried out with WT and hMcm2ΔC (Mcm2-WHD truncated mutant) hMCM2-7. (**c**) Comparison between yeast and human Mcm2 domain organisation. hMcm2ΔC lacks aa826-904, the WHD domain (red). (**d**) Purification of the hMcm5_ΔC_ truncation mutant results in sub-stoichiometric MCM2-7 complexes compared to purified hMCM2-7_WT_. (**e**) Structural comparison of the human and yeast Mcm6-Cdt1 interface and sequence alignment of the corresponding region.

In agreement with studies carried out in budding yeast, the WHDs of hMcm3 and hMcm7 associate via long loops with Orc2 and Orc1 respectively^16, 18^. Unexpectedly, no experimental density was observed in the position occupied by the neighbouring WHD of Mcm4, which was observed in the yeast OCCM (PDB: 5V8F) (Fig. 8a)^16^. Interestingly, in higher eukaryotes the Mcm4-WHD amino acid diverged (Supplementary Fig. 7), hinting that this WHD may have obtained a different role.

We observed clear densities for the WHDs of hMcm2 and hMcm5, which are involved in interactions with Orc subunits. Yeast does not have a Mcm2-WHD, while the yMcm5-WHD was previously not observed in the context of the yOCCM (Fig. 8a).

To assess the role of the hMcm2 WHD in promoting the licensing reaction, we produced the recombinant helicase incorporating an Mcm2-WHD truncation, missing aa826-904 (Fig. 8c) and tested loading efficiency in the pre-RC reaction. Under low salt wash conditions, the signal intensity for hCdc6 was slightly enhanced in reactions employing the Mcm2-WHD mutant when compared to reactions carried out using wild type helicase, indicating reduced Cdc6 release (Fig. 8b, compare lanes 1 and 2). After the high salt wash, we observed with the truncation mutant a reduction in high salt stable MCM2-7 loading (Fig. 8b, compare lanes 3 and 4). Thus, we conclude that the hMcm2-WHD enhances licensing efficiency, consistent with the concept that multiple hORC-MCM2-7 interactions function synergistically during helicase loading. Importantly, when comparing the hOrc1-5-DNA structure with the hOCCM, it is clear that the hCdc6-induced 12.8° rotation of hOrc3 enables the Orc3-Mcm2 contact, as this pushes Orc3 further out and generates proximity to the hMcm2-WHD (Fig. 6c, d, star). Since yeast lacks the entire Mcm2-WHD (Fig. 8c), the evolution of this domain in higher eukaryotes corresponds with enhanced functionality in the licensing reaction, in agreement with our biochemical data. It is also possible that the hMcm2-WHD has additional roles in DNA replication, perhaps serving as a platform for unknown regulators of human DNA replication.

We next attempted to generate a recombinant helicase bearing Mcm5-WHD truncation spanning 664-734 to test in the pre-RC assay. Unfortunately, purification of this mutant MCM2-7 complex resulted in poor Mcm5 retention (Fig. 8), indicating that the Mcm5 WHD contributes to overall complex stability, rendering their analysis in pre-RC formation impossible.

The yMcm6-WHD has a critical role in yOCCM formation, where it functions to regulate DNA insertion and ATP-hydrolysis^7, 17, 69^. Consistent with this, we observed that the hMcm6-WHD is similarly organised as the yMcm6 WHD. It is sandwiched between Orc4, Orc5 and Cdt1 (Fig. 5c, d, Supplementary Fig 5b). Specifically, budding yeast Orc4 contains a long loop that is inserted into the interface of the yMcm6 WHD and its C-terminal AAA+ domain (Supplementary Fig 5c and 5d). However, no functional relevance for this has been observed in yeast^69^. Consistently, this region is absent in hOrc4 (Supplementary Fig 5c). Instead, the critical Mcm6-Cdt1 interaction interface is well conserved between budding yeast and human (Fig. 8e). Here, we observed a close contact between hCdt1 (aa413-424) and hMcm6 (aa756-786). Previously, hMcm6 E757 and L766 and hCdt1 R425, I426, K429 and K433 were identified to be important in pull down assays^62, 70^. However, in this NMR analysis (PDB: 2LE8), the short Cdt1 peptide has substantially shifted^70^ when compared to the hOCCM structure, indicating that structural context is necessary to fully understand the interaction. Thus, the hOCCM structure can now explain how hCdt1 and Mcm6 interact in the context of full-length proteins within the hOCCM complex.

### Structural changes in Orc2 connect ORC auto-inhibition to Mcm3-WHD recruitment

Crucially, the AAA+ domain of hCdc6 stabilizes the hOrc2 WHD in the hOCCM, thus revealing for the first time its active conformation (Fig. 9a, left). In contrast, in hOrc2-5^30^ the hOrc2-WHD was folded inwards blocking DNA interactions (Fig. 9a, centre), something also seen in the *Drosophila* Orc1-6 complex. (Fig. 9a, right) This rotation of the Orc2-WHD defines the autoinhibited state, wherein the ORC complex cannot bind DNA. Consistently, without hCdc6 stabilising the hOrc2-WHD, the flexible hOrc2-WHD was not resolved in the hOrc1-5-DNA structure^30^. Crucially, the hOrc2-WHD conformation observed here is specifically recognised by hMcm3 (Fig. 9b). This data brings to light the critical role hCdc6 plays in positioning the hOrc2-WHD, to activate the ORC complex and facilitate the observed hOrc2-WHD-Mcm3 contacts.

**Fig. 9:**
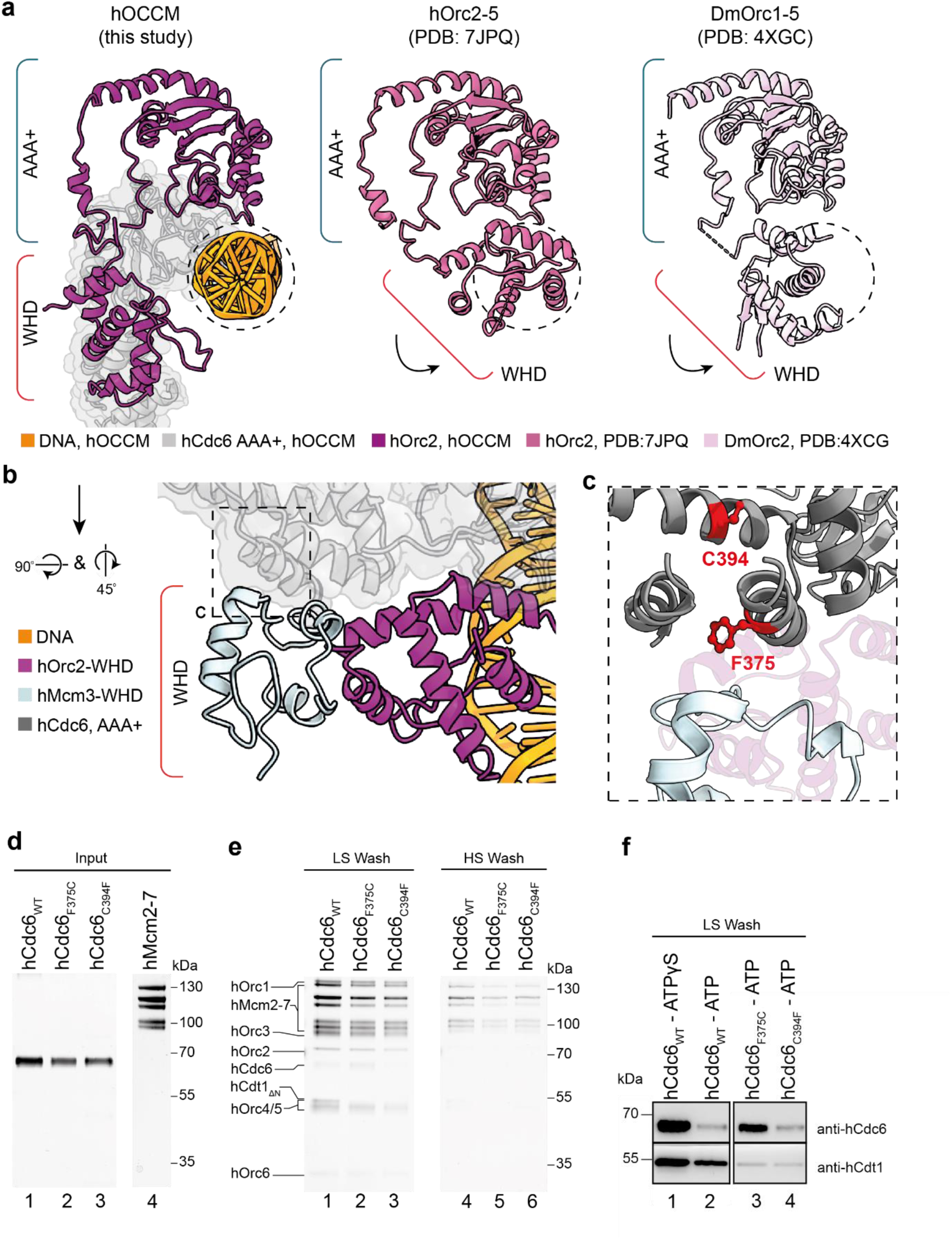
The hOrc2-WHD in the hOCCM is reorganised upon DNA and hCdc6 binding. (**a**) Structure of hOrc2 WHD. The hOrc2 (mauve) is stabilised in the open claw position by the presence of hCdc6 AAA+ domain (light grey), while the closed claw conformations of Orc2 WHD in hOrc2-5 (PDB: 7JPQ) and in DmOrc2 (PDB: 4XCG) not allowing interaction with DNA. (**b**) Structure of hOrc2 WHD (mauve) in the presence of hMcm3 (light blue). (**c**) Localisation of the cosmic mutations in hCdc6. (**d**) Purified WT hCdc6 and F375C and C394F hCdc6 mutants. (**e**) Pre-RC assay under low and high salt wash conditions was carried out with WT hCdc6 and F375C and C394F hCdc6 mutants. (**f**) Corresponding western blot analysis for hCdc6 and hCdt1 in the presence of WT Cdc6 and F375C and C394F hCdc6 mutants. ATPγS arrests complex formation at the hOCCM stage and shows maximal Cdc6 and hCdt1 association.

In budding yeast, it is well established that the yMcm3-yCdc6 interaction initiates pre-RC formation. Our data show that the hCdc6 AAA+ lid and WHD participate in the interaction with hMcm3 (Fig. 5d and 9c). Interestingly, inspection of the <u>C</u>atalogue <u>O</u>f <u>S</u>omatic <u>M</u>utations In <u>C</u>ancer (COSMIC) revealed 83 missense mutations in this region. Structural analysis of these mutations using MutaBind2^71^ highlighted that only few of these mutations are predicted to have a detrimental phenotype. F375C has been found in a lymphoid neoplasm and C394F in adenocarcinoma and non-small cell carcinoma. While hCdc6 F375 is localised directly at the interface with the Mcm3 WHD, hCdc6 C394 is pointing into the inner core of the Cdc6 AAA+ lid domain (Figure 9c). We purified these hCdc6 mutants (Fig. 9d) and asked whether they impact DNA licensing.

Both mutants led to a reproducible reduction in DNA licensing, visible under low and high salt wash conditions (Fig. 9e, lanes 2-3 and 5-6). Specifically, C394F led to diminished Cdt1 and MCM2-7 recruitment, as observed after a low salt wash, and reduced high salt stable MCM2-7 loading, but did not impact on Cdc6 recruitment (Fig. 9f). Due to the introduction of a bulky hydrophobic amino acid into the core of the Mcm3-WHD, it is likely that this mutation is affecting the stability of Mcm3-WHD domain, hence resulting in random unfolding events that reduce activity, but do not block one specific step in complex formation. With F375C, we observed reduced Cdt1 and MCM2-7 recruitment after the low salt wash and reduced high salt stable MCM2-7 loading (Fig. 9f). In contrast, Cdc6 was stabilised (Fig. 9f). Since Cdc6 release is ATP-hydrolysis dependent (Fig. 9f, compare lanes 1 and 2), the data suggest that this mutant affects a step prior to ATP hydrolysis, consistent with a specific defect in Mcm3 recruitment. In summary, our data demonstrate that cancer-associated mutations can impair DNA licensing *in vitro* and that the assay can provide initial insights into their mechanism of action. Consistent with the essential role of Mcm3 in DNA replication and cellular viability, the impact of the mutations was not severe. Moreover, one has to consider that most cancer-associated mutations are heterozygous, thus should be tolerated more easily.

## SUMMARY

Here we have developed a powerful reconstituted system that enables the mechanistic, structural and functional analysis of DNA licensing (Fig. 1). The assay is highly specific (Fig. 1g) and results in the assembly of salt stable hMCM2-7 double-hexamers (Fig. 1 and 3), the final product of the DNA licensing reaction. We show that yeast and human MCM2-7 DNA licensing results in complexes of similar salt stability (Fig. 3). Using the assay, we identify that hOrc6 does not participate in the loading of the first MCM2-7 hexamer (Fig. 1). This is markedly different from budding yeast, where yOrc6 forms an integral part of the complex and is involved in DNA bending during yORC-Cdc6-DNA complex formation^13^. This bending step is critical during DNA licensing, as it generates space for yMCM2-7 recruitment^17^. Future structural work has the potential to reveal if and how hOrc6 independent DNA bending occurs during hORC-Cdc6-DNA complex formation. Interestingly, while yCdt1 forms a stable complex with yMCM2-7, hCdt1 and hMCM2-7 do not interact in solution (Fig. 2). Consistently, hCdt1 is not essential for initial hMCM2-7 recruitment (Fig. 2c). The structure of the hOCCM provided deep insights into the overall complex organisation (Fig. 5). We showed that hCdc6 alters the organisation of hOrc1-5, which supports in turn the establishment of multiple interactions with hMcm-WHDs (Fig. 6). Moreover, our structure-function analysis uncovered how the hMcm2-WHD, which does not exist in yeast, acts in human DNA licensing (Fig. 6 and 8). Our structure provided a snapshot of the active hOrc1-Cdc6 ATPase interface. Interestingly, by analysing a hCdc6 ATPase mutant we discovered that Cdc6 ATPase activity is essential for complex disassembly (Fig. 4). In contrast, yCdc6 ATPase activity has only a role in quality control, e.g. when the complex becomes phosphorylated^7, 18^. Finally, we used the assay to uncover the impact of cancer-associated mutations in hCdc6 and revealed that selected mutants at the hCdc6-Mcm3 interface led to a reduction in DNA licensing activity (Fig. 9). Whether or not this finding is of functional impact to patients is currently unknown, but will be of interest in the future. The established licensing system will be key to generate future mechanistic insight into the human DNA replication, will provide insights into disease causing mutations and is well suited to understand the regulation of the DNA licensing process.

## ACKNOWLEDGEMENTS

We thank members of the Speck lab for critical reading of the manuscript. We thank Audrey Mossler for help with expression and purification of proteins. C.S. was funded by Cancer Research UK (DRCNPG-May21\100006) and the Wellcome Trust (107903/Z/15/Z). We thank the LMS and ICL cryo-EM facilities for support. We thank the LonCEM cryo-EM facility funded by Wellcome Trust (206175/Z/17/Z) and Nora Cronin for support with the data collection.

## AUTHOR CONTRIBUTIONS

J.N.W., V.L. and C.S. conceived the project and designed the experimental approaches. V.L. established the DNA licensing assay and initial protein purification conditions. J.N.W., V.L., L.V.E., S.A., M.P., J.T., A.K.S. and C.M.P. optimised protein purification and purified proteins. J.N.W., V.L., L.V.E., M.P., J.T., S.V.F. and L.R., performed the biochemical assays. J.T. carried out statistical analysis of biochemical assays. M.P. carried out Mass Photometry experiments. J.N.W. and V.L. and characterised protein complexes for cryo-EM structural determination and performed initial sample screenings. J.N.W. prepared cryo-EM samples and performed cryo-EM imaging with support/input from R.A.. J.N.W. processed cryo-EM data with input from A.K.S. and R.A.. A.K.S and J.N.W. built the atomic models with support from R.A.. J.N.W. and C.S. interpreted the structural data. J.N.W prepared figures. J.N.W. and C.S. wrote the manuscript with critical input from all authors. C.S. and V.L. acquired funding for this study.

## COMPETING INTERESTS STATEMENT

The authors have no interests to declare.

## MATERIALS AND METHODS

### Expression and purification of hOrc1-5

hOrc1-5 expression plasmids were generated by gene synthesis (Genscript) based on pESC vectors (Stratagene) for the galactose inducible overexpression in budding yeast; yielding pCS1026, containing ORFs for hOrc5 and hOrc1 as an N-terminal fusion protein with a strep tag, pCS1099 containing the wild-type hOrc2 ORF, and pCS1103, encoding the full-length wild-type hOrc3 and hOrc4 ORFs. The proteins were expressed in *S. cerevisiae* YC658 cells (MATa, lys2::pGAL1 GAL4::LYS2, pep4::HIS3, bar1::hisG derived from W303)^72^ using established conditions^24^. The cells were broken using a SPEXTM freezer-mill. Lysates were resuspended in lysis buffer (50 mM HEPES-NaOH (pH 7.6), 250 mM KCl, 50 mM NaCl, 2 mM DTT, 5 % (w/v) Glycerol, 2 mM MgCl_2_, 0.2 % (w/v) NP-40, 10 mM Β-glyceroPO_4_) supplemented with benzonase (Merck), complete EDTA-free protease inhibitor tablets (Roche) and PhosSTOP phosphatase inhibitor cocktail tablets (Roche). Clarified lysates were then loaded onto a StrepXT column (Cytiva) followed by a wash with lysis buffer, and elution using lysis buffer supplemented with 75 mM biotin. Subsequently, eluted protein was supplemented with 2 mM MgCl_2_, 1 mM ATP, and diluted to 200 mM NaCl for ion exchange chromatography. In some cases, eluates were incubated with SUMOstar protease (LifeSensors) overnight to remove N-terminal StrepTag on hOrc1. Subsequently, protein was bound to a 1 mL HiTrap Heparin column (Cytiva). The column was washed with low salt buffer (50 mM HEPES-NaOH (pH 7.6), 200 mM NaCl, 1.5 mM DTT, 25 mM NaF, 5 % (w/v) Glycerol, 2 mM MgCl_2_, 0.02 % (w/v) NP-40), before being eluted using a salt gradient increasing the buffer to 1M NaCl. Peak fractions were pooled and concentrated for loading onto a 10-30% 400μL glycerol density gradient prepared in 20 mM HEPES-NaOH (pH 7.6), 300 mM NaCl, 1 mM DTT, 2 mM MgCl_2_, 1 mM ATP, and glycerol. Gradients were subjected to ultracentrifugation at 102,000 x g for 16 hours. After manual gradient fractionation, SDS-PAGE and Coomassie staining was used to identify fractions containing stochiometric protein. Finally, target fractions were pooled, concentrated and buffer-exchanged into a final buffer of 20 mM HEPES-NaOH (pH 7.6), 300 mM NaCl, 1 mM DTT, 2 mM MgCl_2_, 1 mM ATP and <10% glycerol, using a 0.5 mL Amicon® Ultra Centrifugal Filter (50 kDa MWCO). Concentrated protein aliquots of ∼3-9 μM were snap-frozen in LN_2_ and stored at –80°C.

### Expression and purification of hOrc6

The hOrc6 expression plasmids were generated by gene synthesis (Genscript) based on pET21a for the IPTG inducible overexpression in bacteria. Plasmid pCS1049, coding for full-length, wild-type hOrc6 with an N-terminal His tag, was transfected into *Escherichia coli* (*E. coli*) Rosetta 1 (DE3) cells (Agilent) and grown in LB media supplemented with ampicillin. Cells were inoculated at an OD_600_ of 0.05 and grown at 37°C to mid-log phase, at which point protein expression was induced by the addition of IPTG. Cells were harvested after 16 h of protein expression at 16°C by centrifugation at 3,500 × g for 10 min. Cell pellets were resuspended in lysis buffer (50 mM HEPES (pH 7.5), 150 mM NaCl, 10mM Imidazole, 5% (v/v) glycerol, 1mM DTT) and lysed by sonication on ice (Branson Digital Sonifier 450/120C). The clarified lysates were then loaded onto a HisTrap Excel column (Cytiva) equilibrated with lysis buffer. The column was washed first using 5 column volume (CV) lysis buffer, then 15 CV of a high salt buffer (lysis buffer supplemented with 1M NaCl) followed by additional 30 CV lysis buffer. Protein was eluted from the column using lysis buffer supplemented with 450 mM imidazole and incubated overnight with PreScission protease (Cytiva) at 4°C to cleave the His-tag. Subsequently, eluted protein was bound to a POROS HQ column equilibrated in buffer A (30mM HEPES-NaOH (pH 7.5), 150mM NaCl, 1mM DTT) for anion exchange chromatography. Protein was eluted across a gradient increasing Buffer B (30 mM HEPES-NaOH (pH 7.5), 1M NaCl, 1mM DTT) to 70% over 15 min. Peak fractions were collected and loaded onto an S75 16/10 column equilibrated in buffer C for size exclusion chromatography. Purified hOrc6 protein was pooled, concentrated in a 30 K Amicon® Ultra-4 concentrator (Millipore), aliquoted, and flash-frozen in LN_2_.

### Expression and purification of wild type and mutant hCdc6

The hCdc6 expression plasmids were generated by gene synthesis (Genscript) based on pGEX6P1 (Cytivia) for the IPTG inducible overexpression in bacteria. Plasmids for wildtype human Cdc6 (pCS536), Cdc6 Walker B mutant E285Q (pCS1339), Cdc6 cosmic mutant F375C (pCS1419) and Cdc6 cosmic mutant C394F (pCS1420) were transfected into *E. coli* BL21 codon + RIL cells (Agilent) and grown in LB media supplemented with ampicillin. Cells were inoculated at an OD_600_ of 0.05 and grown at 37°C to mid-log phase, at which point protein expression was induced by the addition of IPTG. Cells were harvested after 2 h of protein expression at 37°C by centrifugation at 3,500 × g for 10 min. Cell pellets were resuspended in lysis buffer (50 mM HEPES-NaOH (pH 7.6), 250mM KCl, 50mM NaCl, 2mM MgCl_2_, 0.02% (v/v) NP40, 10% (v/v) Glycerol, 2mM DTT) and lysed by sonication (Branson Digital Sonifier 450/120C). Clarified lysates were incubated with GST-Agarose resin (Sigma) for 1 hour at 4°C on a rotator. The resin was washed with 10 CV lysis buffer supplemented with 1mM ATP, 10 CV High Salt buffer (50 mM HEPES-NaOH (pH 7.6), 1 M KCl, 50 mM NaCl, 2 mM MgCl_2_, 0.02% (v/v) NP40, 10% (v/v) Glycerol, 1 mM DTT, 1 mM ATP), and 3 CV buffer C (30mM HEPES-NaOH (pH 7.6), 167mM KCl, 33.2 mM NaCl, 1mM MgCl2, 0.02% (v/v) NP40, 5% (v/v) Glycerol, 1mM DTT, 1mM ATP). Finally, 2 CV Buffer C was added to the resin along with PreScission protease (Cytiva) After O/N cleavage at 4°C the eluted protein was loaded onto a HiTrap SP HP column (Cytiva) equilibrated in buffer C for cation exchange chromatography. Protein was eluted across a gradient increasing buffer D (30mM HEPES-NaOH (p.H. 7.6) 825mM KCl, 165mM NaCl, 1mM DTT, 5 % (w/v) Glycerol, 1mM MgCl_2_, 0.05 % (v/v) NP-40) to 100%. Purified hCdc6 protein was pooled, concentrated in a 30 K Amicon® Ultra-4 concentrator (Millipore), aliquoted, and flash-frozen in liquid nitrogen.

The Walker B mutant hCdc6 was further purified by gel filtration using a Superdex 75 HiLoad column.

The COSMIC hCdc6 mutants were concentrated directly following elution from the affinity resin using 15 mL Amicon® Ultra Centrifugal Filter (30 kDa MWCO) and buffer exchanged into final storage buffer (10mM HEPES pH 7.6, 250mM KCl, 50mM NaCl, 1mM DTT) before being aliquoted and flash frozen without further purification.

### Expression and purification of wild type and mutant hCdt1

The hCdt1 expression plasmids were generated by gene synthesis (Genscript) based on pGEX6P1 (Cytivia) for the IPTG inducible overexpression in bacteria. Plasmids pCS535, coding for full-length Cdt1 fused to an N-terminal GST tag, or pCS1273 coding for N-terminally truncated hCdt1_158-546_ fused at the N-terminus to a GST tag, were transfected into *E. coli* BL21 (DE3) (Agilent) and grown in LB media supplemented with ampicillin. Cells were inoculated at an OD_600_ of 0.05 and grown at 37°C to mid-log phase, at which point protein expression was induced by the addition of IPTG. Cells were harvested after overnight protein expression at 16°C by centrifugation at 3000 × g for 20 min. Cell pellets were resuspended in lysis buffer (50mM HEPES-NaOH (pH 7.6), 250mM NaCl, 2mM DTT, 0.1% (v/v) Triton X-100, 10% (v/v) glycerol) and lysed via sonication (Branson Digital Sonifer 450/120C). The clarified lysate was added to preequilibrated Sepharose Glutathione FastFlow resin (Sigma) and incubated at 4°C for 2 hours. PreScission protease (Cytivia) was added and the solution was incubated overnight at 4°C. The protein was eluted by washing with buffer B (30mM HEPES-NaOH (pH 7.6), 200mM NaCl, 1mM DTT, 0.01% (v/v) Triton X-100, 5% (v/v) glycerol) then was loaded onto a POROS™ HS 20 µm column (ThermoFisher) and eluted using a 200 to 1000 mM NaCl gradient. The protein was concentrated using a 30 K Amicon Ultra-4 concentrator (Millipore) and further purified by gel filtration chromatography on a Superdex 200 Increase 10/300 GL column (Cytiva) equilibrated in final storage buffer (10mM HEPES-NaOH (pH 7.6), 200mM NaCl, 1mM DTT, 0.01% (v/v) Triton X-100, 5% (v/v) glycerol). Protein peak fractions were pooled, concentrated, aliquoted, and flash-frozen in liquid nitrogen.

### Expression and purification of wild type and mutant hMcm2-7

hMcm2-7 complexes were produced in HEK293T cells as described before^73^. The cell mass was resuspended in 25mL lysis buffer (20mM HEPES-NaOH (pH 7.5), 250mM KGlu, 5mM MgCl_2_, 2mM ATP, 10mM imidazole, 0.02% (v/v) NP-40, 5% (v/v) glycerol) containing one SIGMAFAST^TM^ EDTA-free protease inhibitor tablet (Sigma) and Benzonase (Millipore). Cells were lysed via sonication (Branson Digital Sonifer 450/120C). The membrane fraction was removed by centrifugation at 41000 x g for 50 min loaded onto Ni-NTA agarose (Qiagen, #30210) with pre-equilibrated lysis buffer. The resin was washed with 5 CV lysis buffer with protease inhibitor and 5 CV lysis buffer without protease inhibitor. The protein was then eluted using 2 CV elution buffer (20mM HEPES pH 7.5, 250mM KGlu, 5mM MgCl_2_, 2mM ATP, 300mM Imidazole, 0.02% (v/v)NP-40, 5% (v/v) glycerol) before loading onto a StrepTrapXT column (Cytiva) equilibrated with buffer B (20mM HEPES-NaOH (pH 7.5), 250mM KGlu, 5mM MgCl_2_, 2mM ATP, 1mM DTT, 0.02% (v/v) NP-40, 5% (v/v) glycerol). The column was washed with 3 CV buffer B before protein was eluted using 3 CV buffer B supplemented with 50mM biotin. Fractions were visualised with SDS-PAGE, pooled and concentrated using an Amicon Ultra-15 centrifugal concentrator (Millipore, MWCO 50kDa). Purified protein was aliquoted, flash frozen in LN_2_ and stored at −80°C.

### Pulldown assay

The indicated proteins (Orc1-5, Orc6, Cdc6) were mixed in a 1:1:2 ratio in pulldown buffer (15mM HEPES-NaOH, 200mM NaCl, 1mM DTT, 2mM MgCl_2_,1mM ATP, 0.1% (v/v) Triton) in the presence or absence of 90bp yeast ARS1 dsDNA containing the A, B1 and B2 elements. After addition of Mag Strep type 3 XT beads (IBA Lifesciences) and a 10 min incubation at 22°C the beads were washed 3 times. Consequently, the proteins were elected with biotin (2mg/ml). The eluted proteins were separated by SDS-PAGE and silver stained.

### The pre-RC assay

The budding yeast DNA licensing assay was performed as described^7^. For the human DNA licensing assay purified proteins (hOrc1-5, hOrc6, hCdc6, hCdt1, hMcm2-7) at a relative ratio of 1 : 2 : 2 : 1 : 1.5 (respectively) were incubated in pre-RC buffer (25mM HEPES-NaOH (pH 7.5), 250mM KGlu, 1mM DTT, 4mM MgOAc, 0.1% (v/v) Triton-X100, 1mM ATP or ATPγS) for 10 min at 30°C with shaking. Next, 600ng of linear human B2-lamin dsDNA, either 2kbp or 3kbp in length and conjugated to magnetic streptavidin beads via 5’ biotinylation, was added to the reaction which was incubated for an additional 20 min under the same conditions. The magnetic beads are washed with either a low salt buffer (pre-RC buffer) to remove excess unbound protein, or a high salt buffer (pre-RC buffer supplemented with 300mM NaCl) to disrupt hydrophilic interactions and leave only those complexes bound and directly encircling the DNA. Finally, DNA-bound complexes are eluted from the beads by briefly incubating in an elution buffer prepared from pre-RC buffer supplemented with 5mM CaCl_2_ and the endonuclease DNaseI (Thermo Scientific). Eluates were subjected to either SDS-PAGE or used directly to prepare cryo-EM grids.

### Negative stain electron microscopy

#### Grid staining

For all samples, 12 μL eluates from the DNA licensing assay containing ∼100ng of DNA-protein complexes were applied to carbon film 300 mesh Cu grids which had been made hydrophilic by glow discharging using a PELCO easiGlow device. Samples were incubated for 30s before blotting, and subsequently washed twice using pre-RC buffer and once using milli-Q grade water. Grids were then stained twice with 2% uranyl acetate.

#### Data Collection

Data were collected with a Thermo Fischer Talos F200i TEM operated at 200 kV. The microscope was equipped with a Falcon 3EC direct electron detector. EPU software was used for automated acquisition of micrographs collected at x73k magnification using a defocus range from –0.9 to –2.9 μm. Magnification settings resulted in a pixel size of 2.0 Å.pixel^−1^.

#### Image processing

Micrographs were imported into cryoSPARC v4.03. Particles were picked using cryoSPARC own implementation of blob picker, followed by inspect particle picks and filtered using NCC score and local power. Particles were extracted into 200×200 pixel size boxes and subjected to iterative 2D classification. After establishing the parameters for particle picking using a blob picker, we compared the data side by side with data collected during cryo-EM grid screening. For hOCCM, the resulting 2D class averages were used as a template for picking during subsequent high resolution cryo-EM data processing.

### Cryo-EM analysis

#### Sample vitrification

For all samples, ∼100 ng of DNA licensing intermediates were applied to 2nm Quantifoil R2/2 holey carbon support grids which had been glow discharged using a PELCO easiGlow. After 30s, grids were blotted for 1s before being plunged into liquid ethane kept at liquid nitrogen temperature for vitrification, using a TFS Vitrobot Mark IV under 95% humidity.

#### Data Collection

Data were automatically collected with EPU software on a Titan Krios TEM (Thermo Fisher Scientific) operated at 300 keV and equipped with a K3 direct electron detector. The total dose was approximately 40 electrons per Å^2^ for a total of 40 frames per micrograph (Table1). A total of 18,728 micrographs was collected with a defocus range between −0.5 to −2.1 μm. Magnification settings resulted in a pixel size of 1.1 Å.pixel^−1^.

#### Image pre-processing

Original image stacks were summed and corrected for drift- and beam-induced motion using cryoSPARC Patch Motion Correction^63^. The estimation of contrast transfer function (CTF) parameters for each micrograph was performed with PatchCTF^63^.

#### Reconstruction

Image processing was carried out using cryoSPARC v4.03. Micrographs were split into 4 subsets and processed in parallel for particle picking and 2D classification, before being merged for 3D reconstructions. Class averages of hOCCM particles from cryo-grid screening data, resolved using blob picker, were used as templates for another particle picking round and low-pass filtered to 50 Å. Picks were pruned to exclude areas containing thick carbon and extracted into 300×300 pixel size boxes (1.1 Å.pixel^-1^ sampling) giving a total number of 268,243 particles which were included as input for an *Ab initio* reconstruction. High resolution particles were sorted from low resolution particles in multiple rounds of heterogeneous 3D refinement. Finally, 8,730 particles were included in a non-uniform 3D refinement to generate a consensus map with a global resolution reported as 6.09 Å. Half maps, refined maps, sharpened maps, and masks used have been deposited to the EMDB (EMD-9566).

#### Model building

AlphaFold models for human DNA licensing proteins were rigid-body docked into the locally refined maps of the hOCCM using UCSF Chimera^75^. This served as a starting point for molecular model building in COOT^76^, which was combined iteratively with refinement using Namdinator^77^ and Phenix^78^. The molecular model was subjected to geometry minimisation. The molecular model was deposited to the PDB: 8RWV.

### Mass Photometry

Mass Photometry measurements were carried out using a TwoMP Mass Photometer (Refeyn) at room temperature. Samples were added to a six well silicone gasket (GraceBioLabs) on a pre-cleaned sample coverslip (Refeyn). Movies were recorded for 60 seconds in the regular field of view using the AcquireMP software (v. 2023R2, Refeyn Ltd), and analysed using the DiscoverMP software (v. 2023R2, Refeyn Ldt). Prior the measurements, BSA (Sigma, 66kDa monomer, 132kDa dimer) and bovine thyroglobulin (Sigma, 670kDa) were measured to produce a linear mass calibration. The maximum mass error for each calibration was below 5%. Measurements were completed in Mass Photometry buffer (pre-RC buffer without Triton-X100). For the hMCM2-7/hCdt1 and yMCM2-7/yCdt1 measurements, 75nM MCM2-7 was incubated with 150nM Cdt1 (750nM for the 10x hCdt1 measurement) for 20 minutes at 30°C in Mass Photometry Buffer. The instrument was focused using the Droplet Dilution routine in the AcquireMP software with the samples diluted 1 in 10 on the instrument. Measurements were repeated in triplicate. Measurements of the other pre-RC proteins were completed as described above but with the proteins diluted to 75nM and immediately measured.

### Quantification of pre-RC silver stain bands

ImageQuant software (v 9.1, Cytiva) was used to convert silver-stained pre-RC SDS-PAGE gel images to grayscale and measure the density of each MCM2-7 protein band relative to the background of the gel. The mean density of MCM2-7 protein bands for each pre-RC reaction was calculated and compared to that of the control reaction to obtain the relative amount of MCM2-7 loading relative to the control reaction. Mean and standard error values were determined using 4 distinct experimental replicates of pre-RC reaction products visualised onSDS-PAGE gels.

## Notes

### Competing Interest Statement

The authors have declared no competing interest.

## REFERENCES

1. Costa, A. & Diffley, J.F.X. The Initiation of Eukaryotic DNA Replication. Annu Rev Biochem 91, 107–131 (2022).

2. Siddiqui, K., On, K.F. & Diffley, J.F. Regulating DNA replication in eukarya. Cold Spring Harb Perspect Biol 5(2013).

3. Diffley, J.F., Cocker, J.H., Dowell, S.J. & Rowley, A. Two steps in the assembly of complexes at yeast replication origins in vivo. Cell 78, 303–16 (1994).

4. Blow, J.J. & Laskey, R.A. A role for the nuclear envelope in controlling DNA replication within the cell cycle. Nature 332, 546–8 (1988).

5. Ticau, S., Friedman, L.J., Ivica, N.A., Gelles, J. & Bell, S.P. Single-molecule studies of origin licensing reveal mechanisms ensuring bidirectional helicase loading. Cell 161, 513–525 (2015).

6. Riera, A. et al. From structure to mechanism-understanding initiation of DNA replication. Genes Dev 31, 1073–1088 (2017).

7. Fernandez-Cid, A. et al. An ORC/Cdc6/MCM2-7 complex is formed in a multistep reaction to serve as a platform for MCM double-hexamer assembly. Mol Cell 50, 577–88 (2013).

8. Bell, S.P. & Stillman, B. ATP-dependent recognition of eukaryotic origins of DNA replication by a multiprotein complex. Nature 357, 128–34 (1992).

9. Gavin, K.A., Hidaka, M. & Stillman, B. Conserved initiator proteins in eukaryotes. Science 270, 1667-71 (1995).

10. Mendez, J. & Stillman, B. Chromatin association of human origin recognition complex, cdc6, and minichromosome maintenance proteins during the cell cycle: assembly of prereplication complexes in late mitosis. Mol Cell Biol 20, 8602–12 (2000).

11. Liang, C., Weinreich, M. & Stillman, B. ORC and Cdc6p interact and determine the frequency of initiation of DNA replication in the genome. Cell 81, 667–76 (1995).

12. Sun, J. et al. Cdc6-induced conformational changes in ORC bound to origin DNA revealed by cryo-electron microscopy. Structure 20, 534–44 (2012).

13. Feng, X. et al. The structure of ORC-Cdc6 on an origin DNA reveals the mechanism of ORC activation by the replication initiator Cdc6. Nat Commun 12, 3883 (2021).

14. Frigola, J. et al. Cdt1 stabilizes an open MCM ring for helicase loading. Nat Commun 8, 15720 (2017).

15. Zhai, Y. et al. Open-ringed structure of the Cdt1-Mcm2-7 complex as a precursor of the MCM double hexamer. Nat Struct Mol Biol 24, 300–308 (2017).

16. Yuan, Z. et al. Structural basis of Mcm2-7 replicative helicase loading by ORC-Cdc6 and Cdt1. Nat Struct Mol Biol 24, 316–324 (2017).

17. Yuan, Z. et al. Structural mechanism of helicase loading onto replication origin DNA by ORC-Cdc6. Proc Natl Acad Sci U S A 117, 17747–17756 (2020).

18. Frigola, J., Remus, D., Mehanna, A. & Diffley, J.F. ATPase-dependent quality control of DNA replication origin licensing. Nature 495, 339–43 (2013).

19. Ticau, S. et al. Mechanism and timing of Mcm2-7 ring closure during DNA replication origin licensing. Nat Struct Mol Biol 24, 309–315 (2017).

20. Miller, T.C.R., Locke, J., Greiwe, J.F., Diffley, J.F.X. & Costa, A. Mechanism of head-to-head MCM double-hexamer formation revealed by cryo-EM. Nature 575, 704–710 (2019).

21. Zhang, A., Friedman, L.J., Gelles, J. & Bell, S.P. Changing protein-DNA interactions promote ORC binding-site exchange during replication origin licensing. Proc Natl Acad Sci U S A 120, e2305556120 (2023).

22. Gupta, S., Friedman, L.J., Gelles, J. & Bell, S.P. A helicase-tethered ORC flip enables bidirectional helicase loading. Elife 10(2021).

23. Remus, D. et al. Concerted loading of Mcm2-7 double hexamers around DNA during DNA replication origin licensing. Cell 139, 719–30 (2009).

24. Evrin, C. et al. A double-hexameric MCM2-7 complex is loaded onto origin DNA during licensing of eukaryotic DNA replication. Proc Natl Acad Sci U S A 106, 20240–5 (2009).

25. Cheng, J. et al. Structural insight into the assembly and conformational activation of human origin recognition complex. Cell Discov 6, 88 (2020).

26. Tocilj, A. et al. Structure of the active form of human origin recognition complex and its ATPase motor module. Elife 6(2017).

27. Siddiqui, K. & Stillman, B. ATP-dependent assembly of the human origin recognition complex. J Biol Chem 282, 32370–83 (2007).

28. Ranjan, A. & Gossen, M. A structural role for ATP in the formation and stability of the human origin recognition complex. Proc Natl Acad Sci U S A 103, 4864–9 (2006).

29. Vashee, S. et al. Sequence-independent DNA binding and replication initiation by the human origin recognition complex. Genes Dev 17, 1894–908 (2003).

30. Jaremko, M.J., On, K.F., Thomas, D.R., Stillman, B. & Joshua-Tor, L. The dynamic nature of the human origin recognition complex revealed through five cryoEM structures. Elife 9(2020).

31. Liu, S. et al. Structural analysis of human Orc6 protein reveals a homology with transcription factor TFIIB. Proc Natl Acad Sci U S A 108, 7373–8 (2011).

32. Dhar, S.K. & Dutta, A. Identification and characterization of the human ORC6 homolog. J Biol Chem 275, 34983–8 (2000).

33. Prasanth, S.G., Prasanth, K.V. & Stillman, B. Orc6 involved in DNA replication, chromosome segregation, and cytokinesis. Science 297, 1026–31 (2002).

34. Xu, N. et al. Cryo-EM structure of human hexameric MCM2-7 complex. iScience 25, 104976 (2022).

35. Li, J. et al. The human pre-replication complex is an open complex. Cell 186, 98–111 e21 (2023).

36. Li, N. et al. Structure of the origin recognition complex bound to DNA replication origin. Nature 559, 217–222 (2018).

37. Hu, Y. et al. Evolution of DNA replication origin specification and gene silencing mechanisms. Nat Commun 11, 5175 (2020).

38. Lee, C.S.K. et al. Humanizing the yeast origin recognition complex. Nat Commun 12, 33 (2021).

39. Hossain, M., Bhalla, K. & Stillman, B. Multiple, short protein binding motifs in ORC1 and CDC6 control the initiation of DNA replication. Mol Cell 81, 1951–1969 e6 (2021).

40. Hossain, M. & Stillman, B. Meier-Gorlin syndrome mutations disrupt an Orc1 CDK inhibitory domain and cause centrosome reduplication. Genes Dev 26, 1797–810 (2012).

41. Symeonidou, I.E., Taraviras, S. & Lygerou, Z. Control over DNA replication in time and space. FEBS Lett 586, 2803–12 (2012).

42. Cook, J.G. Replication licensing and the DNA damage checkpoint. Front Biosci (Landmark Ed) 14, 5013–30 (2009).

43. McIntosh, D. & Blow, J.J. Dormant origins, the licensing checkpoint, and the response to replicative stresses. Cold Spring Harb Perspect Biol 4(2012).

44. Ge, X.Q., Jackson, D.A. & Blow, J.J. Dormant origins licensed by excess Mcm2-7 are required for human cells to survive replicative stress. Genes Dev 21, 3331–41 (2007).

45. Woodward, A.M. et al. Excess Mcm2-7 license dormant origins of replication that can be used under conditions of replicative stress. J Cell Biol 173, 673–83 (2006).

46. Ibarra, A., Schwob, E. & Mendez, J. Excess MCM proteins protect human cells from replicative stress by licensing backup origins of replication. Proc Natl Acad Sci U S A 105, 8956–61 (2008).

47. Matson, J.P. et al. Rapid DNA replication origin licensing protects stem cell pluripotency. Elife 6(2017).

48. Shreeram, S. & Blow, J.J. The role of the replication licensing system in cell proliferation and cancer. Prog Cell Cycle Res 5, 287–93 (2003).

49. Mei, L. & Cook, J.G. Efficiency and equity in origin licensing to ensure complete DNA replication. Biochem Soc Trans 49, 2133–2141 (2021).

50. Gardner, N.J. et al. The High-Affinity Interaction between ORC and DNA that Is Required for Replication Licensing Is Inhibited by 2-Arylquinolin-4-Amines. Cell Chem Biol 24, 981–992 e4 (2017).

51. Nielsen-Dandoroff, E., Ruegg, M.S.G. & Bicknell, L.S. The expanding genetic and clinical landscape associated with Meier-Gorlin syndrome. Eur J Hum Genet 31, 859–868 (2023).

52. Keskin Karakoyun, H., et al. Evaluation of AlphaFold structure-based protein stability prediction on missense variations in cancer. Front Genet 14, 1052383 (2023).

53. Buel, G.R. & Walters, K.J. Can AlphaFold2 predict the impact of missense mutations on structure? Nat Struct Mol Biol 29, 1–2 (2022).

54. Tate, J.G. et al. COSMIC: the Catalogue Of Somatic Mutations In Cancer. Nucleic Acids Res 47, D941–D947 (2019).

55. Speck, C. & Stillman, B. Cdc6 ATPase activity regulates ORC x Cdc6 stability and the selection of specific DNA sequences as origins of DNA replication. J Biol Chem 282, 11705–14 (2007).

56. Bleichert, F., Leitner, A., Aebersold, R., Botchan, M.R. & Berger, J.M. Conformational control and DNA-binding mechanism of the metazoan origin recognition complex. Proc Natl Acad Sci U S A 115, E5906–E5915 (2018).

57. Dhar, S.K. et al. Replication from oriP of Epstein-Barr virus requires human ORC and is inhibited by geminin. Cell 106, 287–96 (2001).

58. Hesketh, E.L., Knight, J.R., Wilson, R.H., Chong, J.P. & Coverley, D. Transient association of MCM complex proteins with the nuclear matrix during initiation of mammalian DNA replication. Cell Cycle 14, 333–41 (2015).

59. Donovan, S., Harwood, J., Drury, L.S. & Diffley, J.F. Cdc6p-dependent loading of Mcm proteins onto pre-replicative chromatin in budding yeast. Proc Natl Acad Sci U S A 94, 5611–6 (1997).

60. Samel, S.A. et al. A unique DNA entry gate serves for regulated loading of the eukaryotic replicative helicase MCM2-7 onto DNA. Genes Dev 28, 1653–66 (2014).

61. Li, N. et al. Structure of the eukaryotic MCM complex at 3.8 A. Nature 524, 186–91 (2015).

62. Wei, Z. et al. Characterization and structure determination of the Cdt1 binding domain of human minichromosome maintenance (Mcm) 6. J Biol Chem 285, 12469–73 (2010).

63. Punjani, A., Rubinstein, J.L., Fleet, D.J. & Brubaker, M.A. cryoSPARC: algorithms for rapid unsupervised cryo-EM structure determination. Nat Methods 14, 290–296 (2017).

64. Bleichert, F., Botchan, M.R. & Berger, J.M. Crystal structure of the eukaryotic origin recognition complex. Nature 519, 321–6 (2015).

65. Lim, Y. et al. In silico protein interaction screening uncovers DONSON’s role in replication initiation. Science 381, eadi3448 (2023).

66. Neuwald, A.F. Hypothesis: bacterial clamp loader ATPase activation through DNA-dependent repositioning of the catalytic base and of a trans-acting catalytic threonine. Nucleic Acids Res 34, 5280–90 (2006).

67. Coster, G., Frigola, J., Beuron, F., Morris, E.P. & Diffley, J.F. Origin licensing requires ATP binding and hydrolysis by the MCM replicative helicase. Mol Cell 55, 666–77 (2014).

68. Herbig, U., Marlar, C.A. & Fanning, E. The Cdc6 nucleotide-binding site regulates its activity in DNA replication in human cells. Mol Biol Cell 10, 2631–45 (1999).

69. Guerrero-Puigdevall, M., Fernandez-Fuentes, N. & Frigola, J. Stabilisation of half MCM ring by Cdt1 during DNA insertion. Nat Commun 12, 1746 (2021).

70. Liu, C. et al. Structural insights into the Cdt1-mediated MCM2-7 chromatin loading. Nucleic Acids Res 40, 3208-17 (2012).

71. Zhang, N. et al. MutaBind2: Predicting the Impacts of Single and Multiple Mutations on Protein-Protein Interactions. iScience 23, 100939 (2020).

72. Matheson, K., Parsons, L. & Gammie, A. Whole-Genome Sequence and Variant Analysis of W303, a Widely-Used Strain of Saccharomyces cerevisiae. G3 (Bethesda) 7, 2219–2226 (2017).

73. Rzechorzek, N.J., Hardwick, S.W., Jatikusumo, V.A., Chirgadze, D.Y. & Pellegrini, L. CryoEM structures of human CMG-ATPgammaS-DNA and CMG-AND-1 complexes. Nucleic Acids Res 48, 6980–6995 (2020).

74. Pettersen, E.F. et al. UCSF Chimera--a visualization system for exploratory research and analysis. J Comput Chem 25, 1605–12 (2004).

75. Emsley, P., Lohkamp, B., Scott, W.G. & Cowtan, K. Features and development of Coot. Acta Crystallogr D Biol Crystallogr 66, 486–501 (2010).

76. Kidmose, R.T. et al. Namdinator - automatic molecular dynamics flexible fitting of structural models into cryo-EM and crystallography experimental maps. IUCrJ 6, 526–531 (2019).

77. Liebschner, D. et al. Macromolecular structure determination using X-rays, neutrons and electrons: recent developments in Phenix. Acta Crystallogr D Struct Biol 75, 861–877 (2019).

