## Supplemental Files for "Reconstitution of human DNA licensing and the structural and functional analysis of key intermediates"

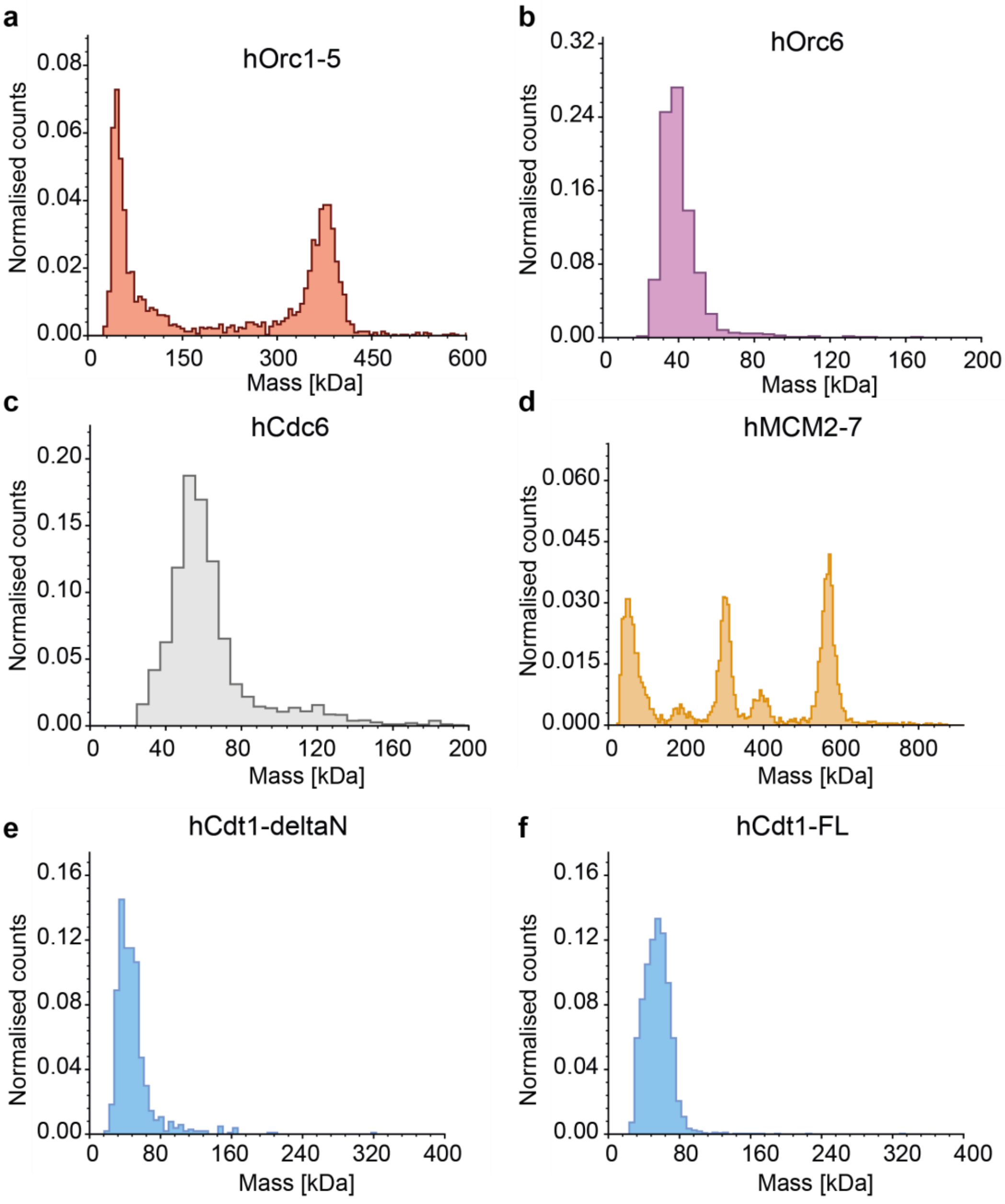

**Supplementary Fig. 1**: **Mass Photometry histograms of purified proteins used in the pre-RC assay**. (**a**) hOrc1-5, (**b**) hOrc6, (**c**) hCdc6, (**d**) hMCM2-7 (**e**) hCdt1 ΔN truncation mutant and (**f**) full length hCdt1.

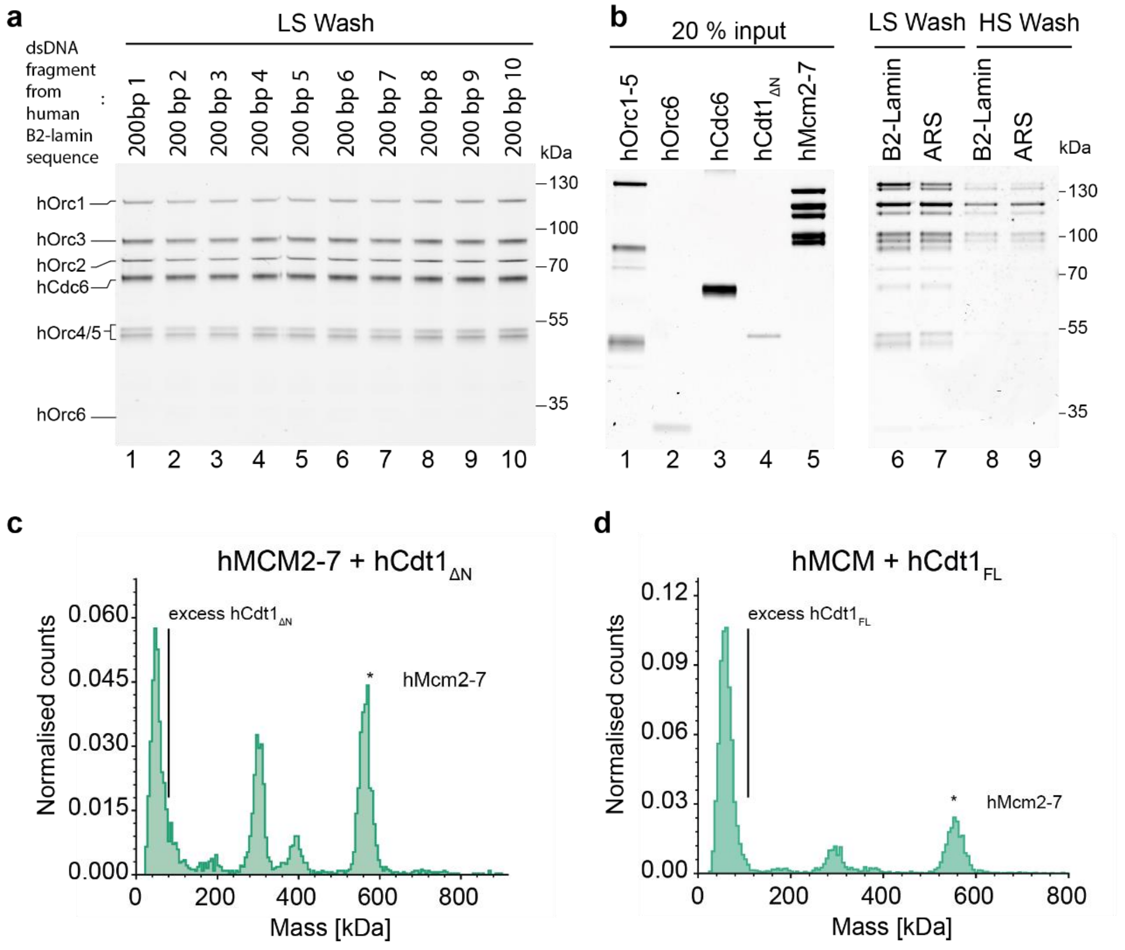

**Supplementary Fig. 2**: **DNA sequence analysis of DNA licensing and interaction analysis of Cdt1-MCM2-7**. (**a**) Pre-RC like assay, hOrc1-6 and hCdc6 were incubated with 200bp dsDNA fragments along the length of the human B2-lamin sequence and washed under low salt wash conditions. (**b**) Pre-RC assay under low and high salt wash conditions was carried out with hB2-Lamin and yARS1 DNA. Mass photometry analysis of the ability of (**c**) hMCM2-7 and hCdt1$_{\Delta N}$ and (**d**) hCdt1$_{FL}$ to interact. This was carried out in solution with a 2-fold excess of hCdt1.

**Supplementary Table 1 Summary of Cryo-EM Data Collection and Model Refinement**

| | *hOCCM* |
|---|---|
| **Data Collection/Processing** | |
| Voltage (kV) | 300 |
| Magnification | 81,000 |
| Defocus range (µm) | -0.5 to -2.1 (0.2) |
| Symmetry imposed | C1 |
| Total electron dose (e-/ A^2) | 40 |
| Exposure Time (s) | 3.0 |
| Number of micrographs | 18,728 |
| Number of frames/micrograph | 40 |
| Initial Particle Number | 334,603 |
| Final Particle Number | 8,730 |
| Resolution (masked, Å) | 6.09 |
| FSC threshold | 0.143 |
| **Refinement** | |
| Model composition | |
| Protein Residues | 6369 |
| Nucleotides | 79 |
| Ligand | 0 |
| RMS Deviations | |
| Bond Lengths (Å) | 0.24 |
| Bond Angels (degree) | 0.49 |
| Ramachandran | |
| Favoured (%) | 87 |
| Allowed (%) | 12 |
| Outlier (%) | 2 |
| MolProbity Score | 5.67 |

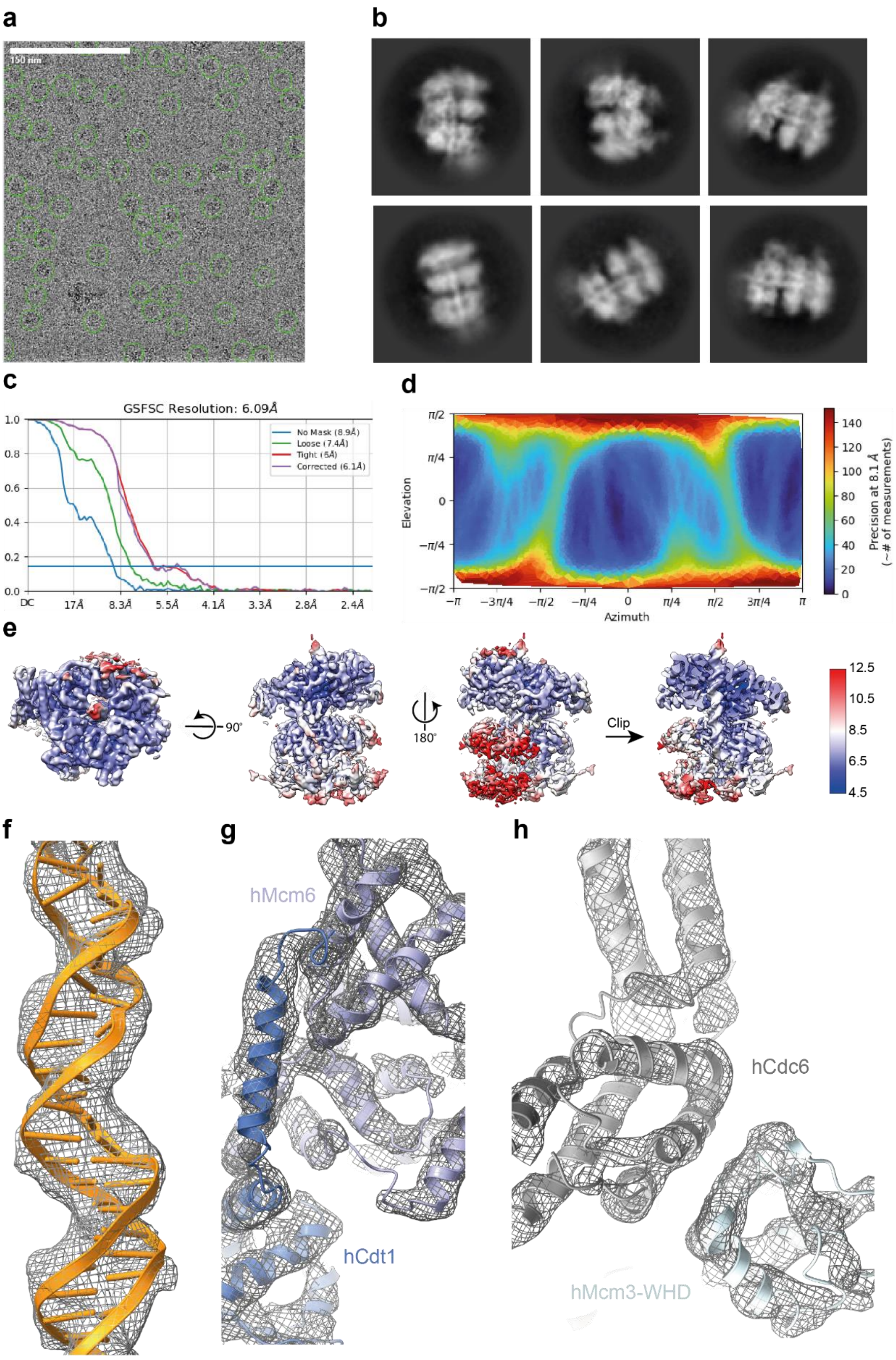

**Supplementary Fig. 3: Quality of the EM data**. (**a**) A typical micrograph was taken on a Titan Krios TEM operated at 300 kV and a K3 direct electron detector. Picked complexes are circled in green. Scale bar is 150 nm. (**b**) Typical class averages show the complex with DNA. (**c**) **P**lot reporting 6.09 Å resolution. (**d**) Angular distribution of the particles used in the final 3D map. (**e**) Local resolution representation of the complex. (**f**) Density map in the area of DNA with the fitted model. (**g**) Density map in the area of the hCdt1-Mcm6 interaction with the fitted model. (**h**) Density map in the area of the hcdc6-Mcm3 interaction with the fitted model.

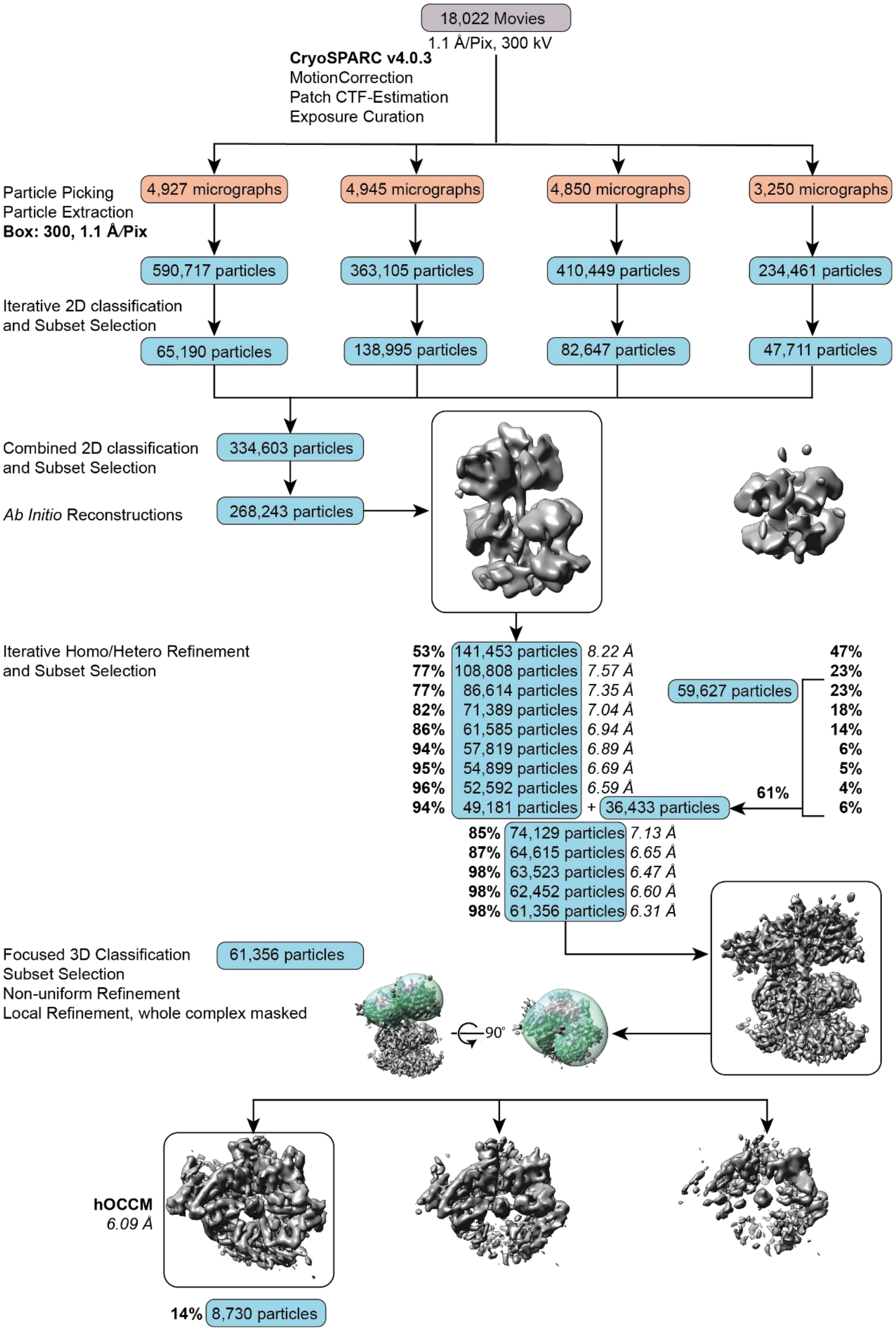

**Supplementary Fig. 4: Image processing workflow.** Processing workflow for hOCCM using cryoSPARC v4.03.

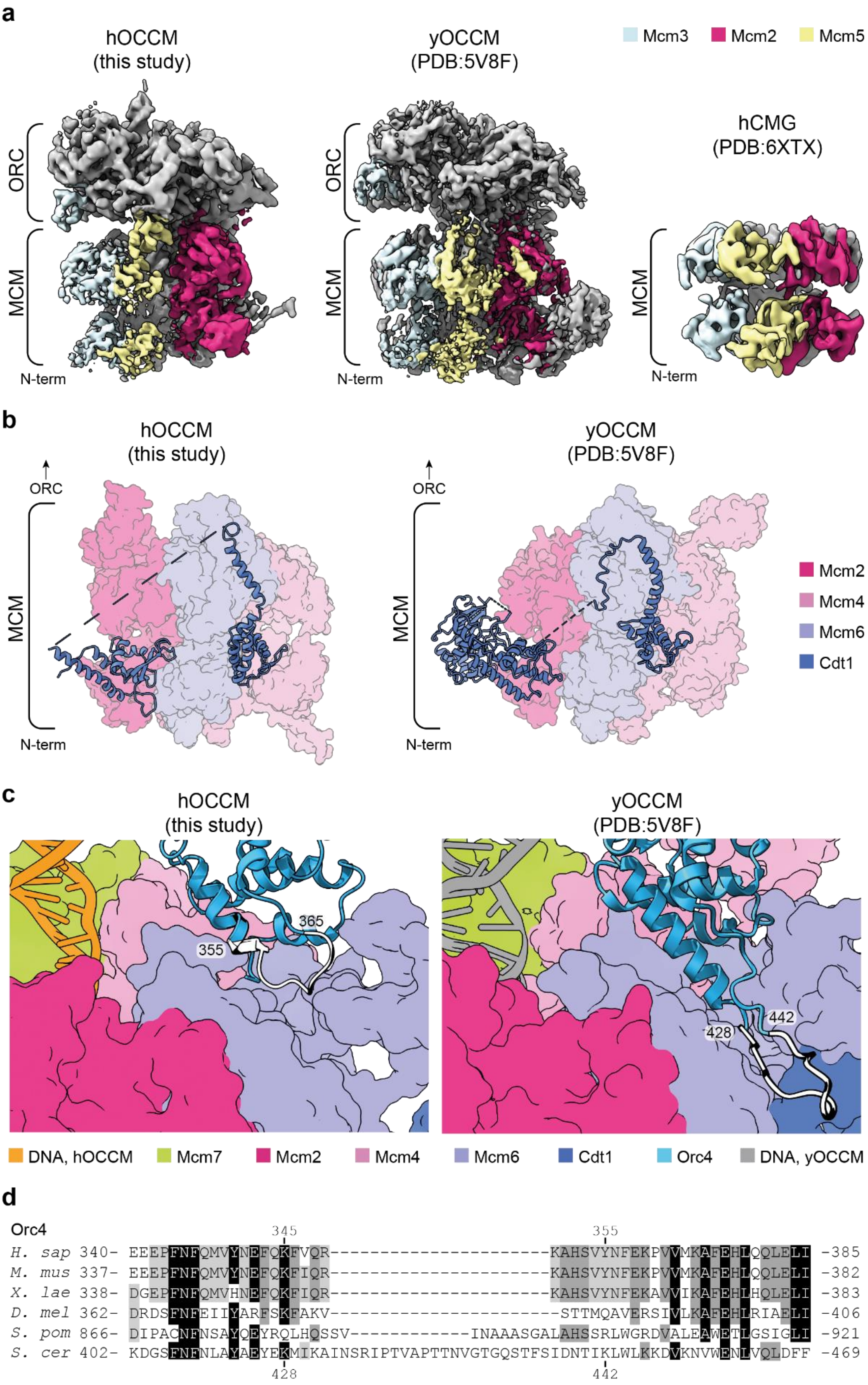

**Supplementary Figure 5: Comparisons between the hOCCM and yOCCM.** (**a**) hOCCM map (left) showing the flexibility of Mcm5 at the 2/5 gate compared to the experimental map for the yOCCM (PDB: 5V8f, centre) and experimental density for the hCMG (PDB: 6XTY, right). All maps are displayed at similar contour levels and have been coloured to highlight experimental density corresponding to Mcm2 (dark pink), Mcm3 (blue), and Mcm5 (yellow). (**b**) Molecular models depicting conservation of the binding position for Cdt1 (blue) bound to Mcm2 (dark pink) Mcm6 (purple) and Mcm4 (light pink), with MCM complexes displayed as surfaces in the hOCCM (left) and yOCCM (PDB: 5V8F, right). (**c**) Orc4 loop position relative to hMcm6 WHD in hOCCM compared to yOCCM. (**d**) Protein sequence alignment for Orc4 loop region

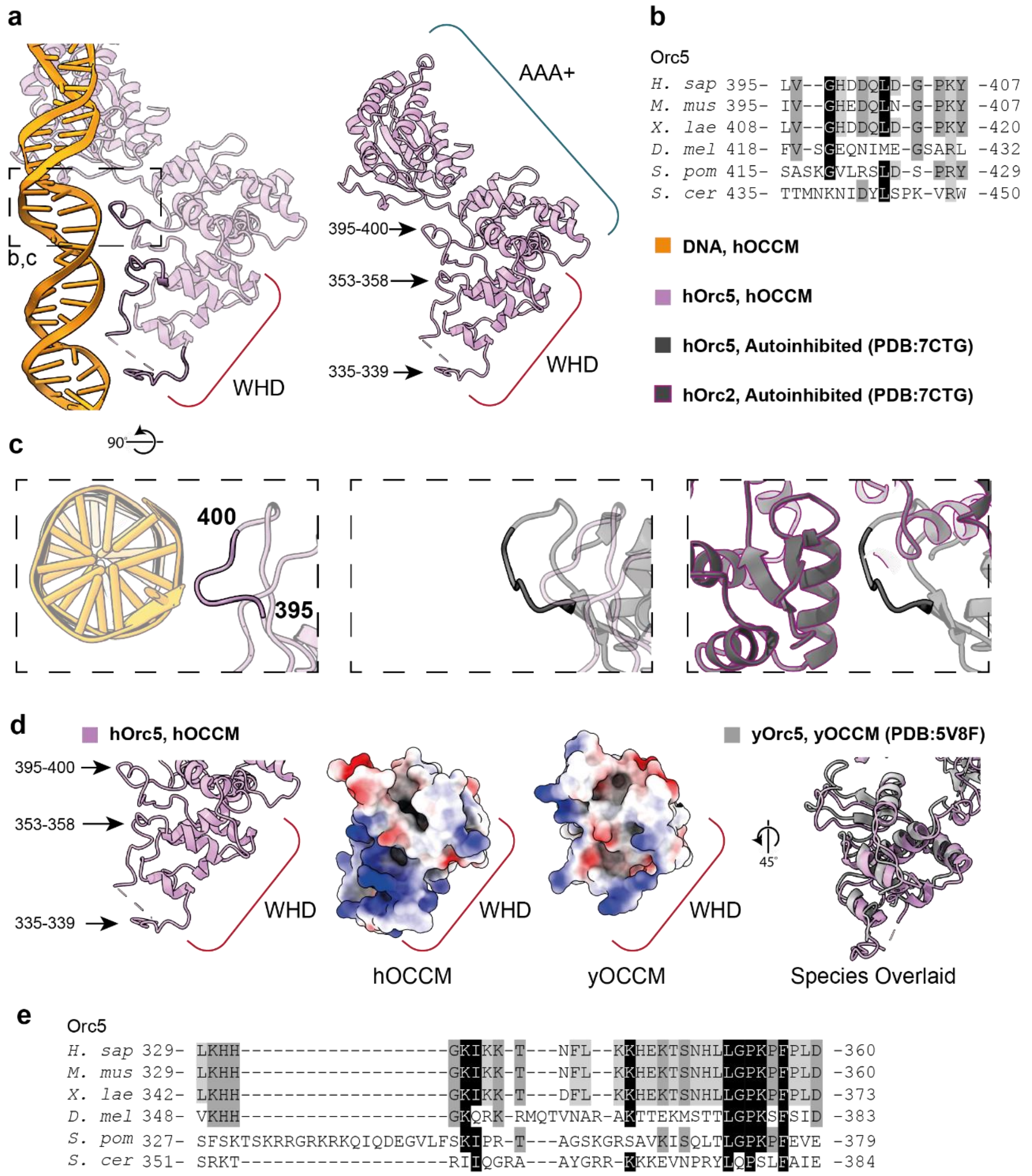

**Supplementary Fig. 6: hOrc5 makes multiple DNA backbone contacts.** (**a**) The hOrc5-WHD makes three contacts with the phosphate backbone. (**b**) Sequence alignment of the hOrc5 aa395-400 DNA contact. (**c**) hOrc5 aa394-400 makes contact with DNA, but in the autoinhibited state, the same region is interacting with hOrc2. (**d**) Comparison of the hOrc5-WHD and the yeast counterpart. (**e**) Sequence alignment of hOrc5 aa329-360, highlighting a conserved sequence in higher eukaryotes.

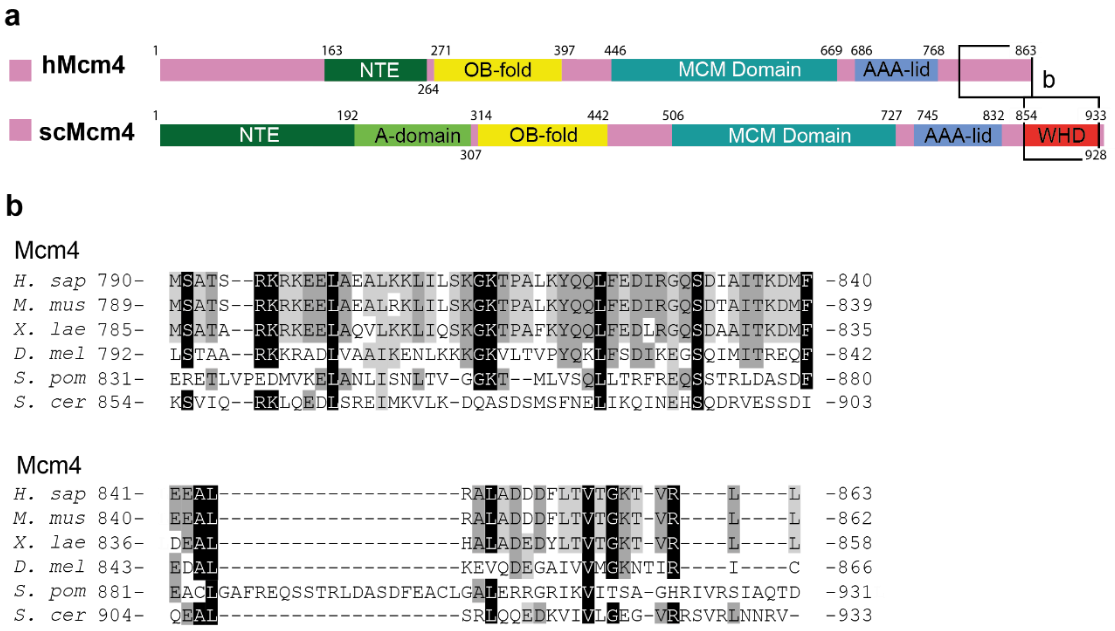

**Supplementary Fig. 7: C-terminal regions of Mcm4 diverge.** (**a**) Side-by-side comparison of the domain organisation of Mcm4 in human (top) and yeast (bottom). (**b**) Alignment of the C-terminus of Mcm4 highlighting its divergence during evolution.

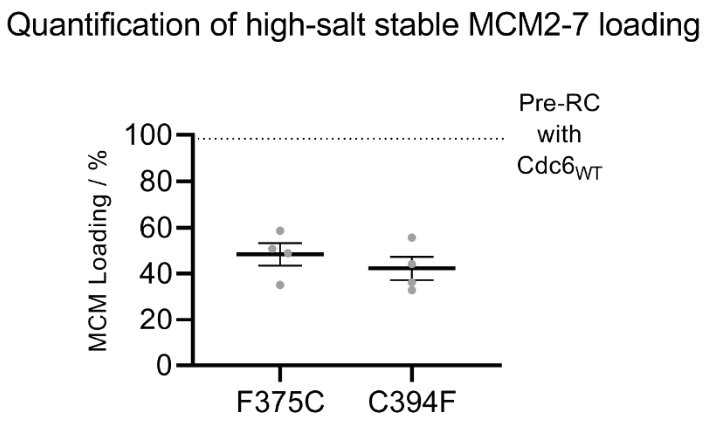

**Supplementary Fig.8: Cosmic mutations in hCdc6 impact high salt-stable MCM2-7 loading.** Quantification of MCM2-7 loading efficiency showing the comparison of WT hCdc6 and two hCdc6 Cosmic mutants from n=4 distinct pre-RC assays.
